# ESYT1 tethers the endoplasmic reticulum to mitochondria and is required for mitochondrial lipid and calcium homeostasis

**DOI:** 10.1101/2022.11.14.516495

**Authors:** Alexandre Janer, Jordan L. Morris, Michiel Krols, Hana Antonicka, Mari J. Aaltonen, Zhen-Yuan Lin, Anne-Claude Gingras, Julien Prudent, Eric A. Shoubridge

## Abstract

Mitochondria interact with the endoplasmic reticulum (ER) at structurally and functionally specialized membrane contact sites known as mitochondria-ER contact sites (MERCs). MERCs are crucial for a myriad of physiological functions including lipid synthesis and transport, and calcium signaling. Alterations in the structure, composition or regulation of MERCs contribute to the aetiology of many pathologies including neurodegenerative and metabolic diseases. The proteins mediating the formation of MERCs have been extensively studied in yeast, where the ER-mitochondria encounter structure (ERMES) complex mediates the transport of lipids between the ER and mitochondria via three lipid binding SMP-domain proteins. However, none of the SMP proteins of the ERMES complex have orthologues in mammals suggesting that alternate pathways have evolved in metazoans. Combining proximity labelling (BioID), confocal microscopy and subcellular fractionation, we found that the ER resident SMP-domain containing protein ESYT1 was enriched at MERCs, where it forms a complex with the outer mitochondrial membrane protein SYNJ2BP. The deletion of ESYT1 or SYNJ2BP reduced the number and length of MERCs, indicating that the ESYT1-SYN2JBP complex plays a role in tethering ER and mitochondria. Loss of this complex impaired ER to mitochondria calcium flux and provoked a significant alteration of the mitochondrial lipidome, most prominently a reduction of cardiolipins and phosphatidylethanolamines. Both phenotypes were rescued by re-expression of wild-type ESYT1 as well as an artificial mitochondria-ER tether. Together, these results reveal a novel function of ESYT1 in mitochondrial and cellular homeostasis through its role in the regulation of MERCs.

## INTRODUCTION

Mitochondria interact with several membrane-delimited organelles within the cell, including the endoplasmic reticulum (ER), lysosomes, peroxisomes and trans-Golgi network vesicles (Tabara, Morris et al. 2021). Mitochondria-ER contact sites (MERCs), also called MAMs (mitochondria-associated membranes) when studied at a biochemical level, are the best characterized class of membrane contact sites (MCSs) and represent the close apposition of the outer mitochondrial membrane with the ER membrane (Giacomello and Pellegrini 2016). MERCs are functionally and structurally specialized subcellular domains that form a signaling platform allowing lipid synthesis and transport, calcium signalling, apoptosis regulation, mitochondrial division, and autophagosome formation (Herrera-Cruz and Simmen 2017, Giacomello, Pyakurel et al. 2020). MERCs have also been shown to be involved in several critical cellular pathways such as metabolic regulation in diabetes (Rieusset 2017), inflammation (Missiroli, Patergnani et al. 2018), the immune response (Martinvalet 2018) and senescence (Janikiewicz, Szymanski et al. 2018). Alterations in these structures have also been linked to the onset of neurodegenerative diseases including Alzheimer’s Disease, Parkinson’s Disease and Amyotrophic Lateral Sclerosis (Vallese, Barazzuol et al. 2020), and aging (Janikiewicz, Szymanski et al. 2018).

The proteins that mediate the formation of MERCs have been extensively studied in the yeast *Saccharomyces cerevisiae*, where a 4-subunit ER-mitochondria encounter structure (ERMES) is required to tether the two organelles and mediate lipid transport from the ER to mitochondria (Kornmann, Currie et al. 2009, Kojima, Endo et al. 2016) via the lipid binding SMP-domains (synaptotagmin-like mitochondrial and lipid-binding protein) present in three subunits of the complex (Kopec, Alva et al. 2010, AhYoung, Jiang et al. 2015).

Mitochondria synthesize cardiolipin (CL) and phosphatidylethanolamine (PE) on the inner membrane, and these lipids are essential for mitochondrial function (Steenbergen, Nanowski et al. 2005, Funai, Summers et al. 2020). CL is produced via a multienzymatic cascade and PE is synthesized by phosphatidylserine decarboxylase PISD1; however, their synthesis depends on the ER for the supply of the precursor lipids phosphatidic acid (PA) and phosphatidylserine (PS) (Funai, Summers et al. 2020). Lipid synthesis activity at MAMs was the first biochemical process reported at a MCS in mammals (Vance 1990); however, a detailed mechanism of lipid transport between ER and mitochondria in mammals remains elusive as none of the 3 SMP domain proteins in the ERMES complex has orthologues in mammals.

All SMP domain containing proteins are present at MCSs, where they are thought to facilitate non-vesicular transport of lipids between lipid bilayers (Jeyasimman and Saheki 2020). In mammals, the ER anchored extended synaptotagmin (ESYTs) proteins are the best characterized (Saheki and De Camilli 2017). ESYT1, ESYT2 and ESYT3 tether the ER to the plasma membrane (PM), potentially transferring lipids (Bian, Saheki et al. 2018). More specifically, ESYT1 has been shown to play a role in Ca^2+^-dependent lipid transfer at ER-PM contacts, which requires its docking with PIP(4,5)P_2_ in the plasma membrane (Giordano, Saheki et al. 2013, Reinisch and De Camilli 2015) (Bian, Saheki et al. 2018, Ge, Bian et al. 2022). It also tethers ER to peroxisomes by a similar mechanism facilitating the transport of cholesterol (Xiao, Luo et al. 2019), raising the possibility that ESYT1 could also tether ER to mitochondria to promote lipid transfer.

In this study, we used the proximity mapping tool BioID to identify and characterize SMP domain proteins that might be involved in ER-mitochondrial lipid transport in humans. We showed that ESYT1 is enriched at MERCs, where it forms a complex with the outer mitochondrial membrane (OMM) protein SYNJ2BP. Depletion of the ESYT1-SYNJ2BP complex impairs mitochondrial calcium uptake capacity and provokes a reduction of essential mitochondrial lipids, demonstrating its essential function in cellular and mitochondrial homeostasis.

## RESULTS

### Proximity labelling analysis of SMP domain proteins in human

We have recently established that the proximity of proteins localized on two different membrane bound organelles can be detected by the proximity mapping tool BioID (Antonicka, Lin et al. 2020, Go, Knight et al. 2021). To identify potential human SMP domain-containing proteins involved in the regulation of MCSs and lipid transport between ER and mitochondria, we selected several ER-resident human SMP domaincontaining proteins as baits (PDZD8, TEX2, ESYT2 and ESYT1). We generated stable inducible Flp-In T-REx 293 cell lines overexpressing each protein fused with BirA* (Figure S1A) to identify their proximity interactomes and potential partners localized on the outer mitochondrial membrane.

BioID analysis of the selected SMP domain containing proteins (Table S1) revealed, as expected, an enriched ER environment with the majority of their proximity interactors being ER membrane proteins, involved in organelle organization, transport, lipid biosynthesis and regulation of protein metabolic processes: 35 out of 41 preys shared among all four baits were ER proteins (Figure S1B, Table S1). In addition, two preys common to all four baits, ALDH3A2 and FKBP8, had been proposed to dually localize to mitochondria and ER (Shirane and Nakayama 2003, Rath, Sharma et al. 2021, Zeng, Li et al. 2021). Finally, each bait detected numerous unique proximity interactors (Figure S1B).

PDZD8 was previously shown to partially localize at MERCs and tether the two organelles (Hirabayashi, Kwon et al. 2017), but its interacting partner on the OMM remains unknown. Due to its capacity to regulate MERCs, the absence of PDZD8 led to decreased mitochondrial calcium uptake capacity upon ER stimulation (Hirabayashi, Kwon et al. 2017). Moreover, it was sufficient to rescue locomotor defects in a fly model of Alzheimer’s disease expressing Amyloid β_42_ (Hewitt, Miller-Fleming et al. 2022). PDZD8 was later described to interact with RAB7 and ZFYVE27 (Protrudin) to establish a three-way membrane contact sites between the ER, late endosomes and mitochondria and to mediate lipid transfer required for late endosome maturation (Elbaz-Alon, Guo et al. 2020, Shirane, Wada et al. 2020, Khan, Chen et al. 2021, Gao, Xiong et al. 2022). Mass spectrometry results obtained with either the N-, or C-terminal PDZD8-BirA* fusion proteins confirmed the proximity interaction with ZFYVE27 but failed to identify any OMM-localized partner (Table S1).

TEX2 is still uncharacterized in mammals; however, its yeast ortholog Nvj2 localizes to ER-vacuole (lysosome-like organelle) contact sites at steady state. Upon ER stress or ceramide overproduction, it translocases to ER-Golgi contacts to facilitate the non-vesicular transport of ceramide from the ER to the Golgi, counteracting ceramide toxicity (Liu, Choudhary et al. 2017). Consistent with the role of Nvj2 in yeast, we identified 12 proteins belonging to the ER-Golgi vesicle-mediated transport pathway in the N-ter TEX2 proximity interactome (Table S1, in green); however, as with PDZD8 we did not identify any OMM proximity interactor.

In contrast to ESYT2 that constitutively tethers ER to the PM and is localized in the cortical ER, the interaction of ESYT1 with the PM is activated by Ca^2+^ binding. The proportion of ESYT1 present throughout the ER or concentrated at ER-PM contacts is controlled by cytosolic Ca^2+^ (Chang, Hsieh et al. 2013, Giordano, Saheki et al. 2013, Idevall-Hagren, Lu et al. 2015). As ESYT members could form heteromeric complexes, ESYT-dependent ER-PM contacts are regulated by both cytosolic Ca^2+^ and the specific phospholipid PI(4,5)P_2_ at the PM (Fernandez-Busnadiego, Saheki et al. 2015). In both N- and C-terminal ESYT1-BirA* experiments (Table S1), we confirmed the interaction with its known partner ESYT2. Importantly, we also found a unique specific proximity interaction with the OMM protein SYNJ2BP (OMP25) (Figure S1C). This interaction was previously noted but never further investigated (Christianson, Olzmann et al. 2011, Hung, Lam et al. 2017). Significantly, ESYT2 BioID analysis also identified ESYT1 (Table S1) as its main proximity interactor but failed to identify SYNJ2BP, suggesting that ESYT1 may form a specific complex with SYNJ2BP at MERCs independent of its interaction with ESYT2 at ER-PM contacts.

These data prompted us to perform a BioID analysis using SYNJ2BP as a bait (Table S1) and we observed a strong enrichment of ESYT1, confirming the proximity interaction of the two partners. SYNJ2BP was shown to interact with another ER-localized protein RRBP1 to regulate the formation of MERCs (Hung, Lam et al. 2017), and as expected we also identified RRBP1 as a prey. Hung et al. also described SYNJ2BP to interact with the multi aminoacyl tRNA synthetase complex (Mirande 2017), an interaction confirmed in our SYNJ2BP BioID results, further validating the specificity of our BioID results.

Thus, of the four SMP domain-containing proteins we profiled only ESYT1 emerged as having a specific proximity interacting partner on the OMM, SYNJ2BP, suggesting it could play a role in MERCs function.

### ESYT1 localizes at mitochondria-ER contact sites

To test whether ESYT1 could play a role at MERCs, we first studied its intracellular localization by immunofluorescence and confocal microscopy (Figure 1A). In human fibroblasts stably overexpressing SEC61B-mCherry as an ER marker (green) and stained for PRDX3 as a mitochondrial marker (cyan), endogenous ESYT1 (magenta) specifically localized along the ER network forming puncta, especially on ER tubules (which function in lipid and hormone synthesis) rather than on the perinuclear sheets (which function in protein synthesis) (Schwarz and Blower 2016). Line scans of fluorescence intensities showed that the focal accumulations of endogenous ESYT1 along the ER network partially colocalized with mitochondria (Figure 1B).

**Figure 1.**
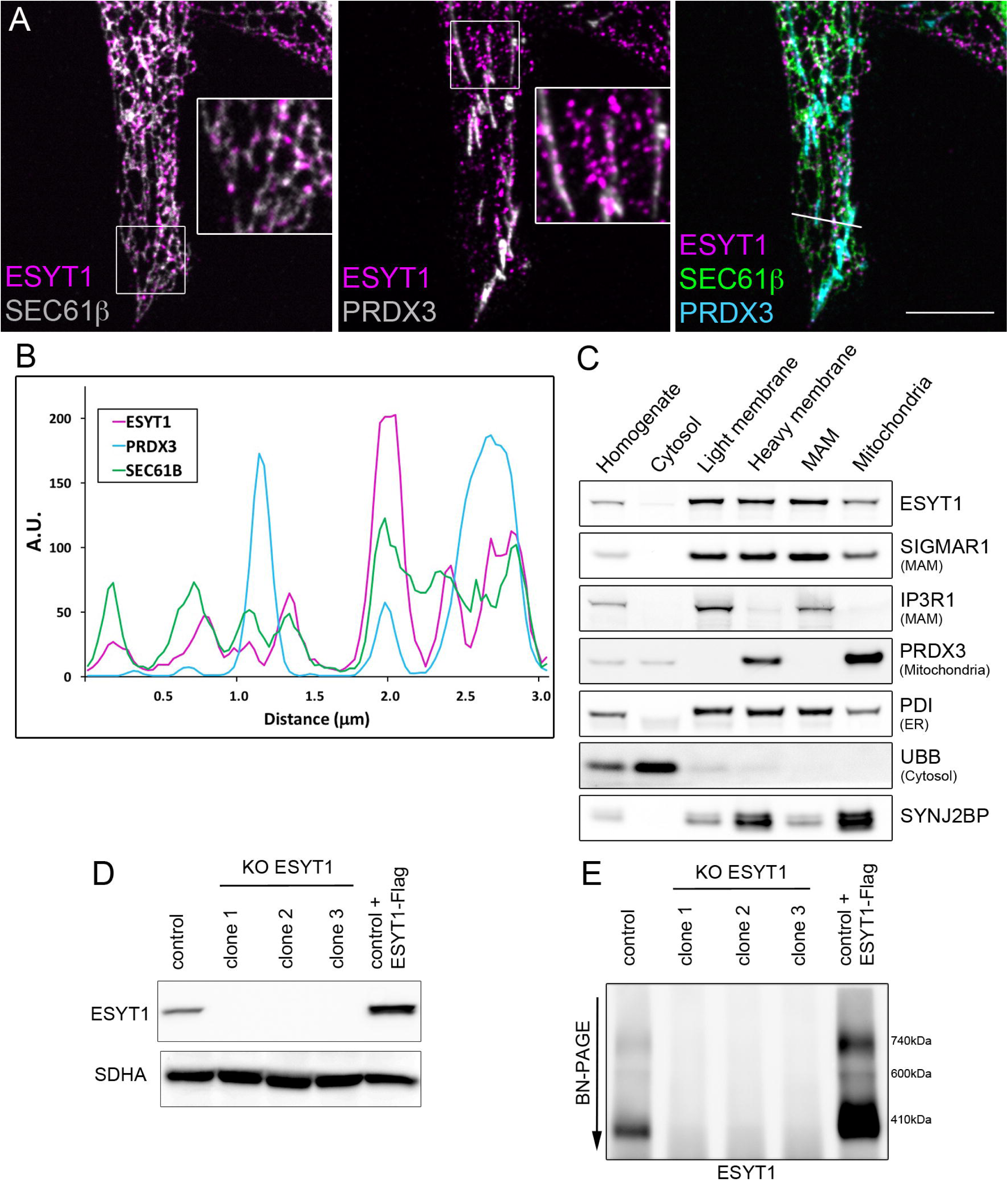
ESYT1 localizes at mitochondria-ER contact sites. A. Confocal microscopy images of endogenous ESYT1 localization (magenta) in human fibroblasts stably overexpressing SEC61B-mCherry as an ER marker (green). Staining for endogenous PRDX3 serves as a mitochondrial marker (cyan). Scale bar=5μm. B. Line scan of fluorescence intensities demonstrating focal accumulations of endogenous ESYT1 along the ER network that partially colocalize with mitochondria (A.U.=arbitrary units) C. Subcellular localization of endogenous ESYT1 and SYNJ2BP. Mouse liver was fractionated, and the fractions were analyzed by SDS-PAGE and immunoblotting. SIGMAR1 and IP3R1 are MAM markers, PRDX3 is a mitochondrial marker, PDI an ER marker and UBB a cytosol marker. D. ESYT1 protein levels in control human fibroblast, three individual clones of ESYT1 knock-out fibroblasts and fibroblasts overexpressing ESYT1-3xFLAG. Whole cell lysates were analyzed by SDS-PAGE and immunoblotting. SDHA was used as a loading control. E. Characterization of the ESYT1 complexes. Heavy membrane fractions were isolated from control human fibroblasts, ESYT1 knock-out fibroblasts and fibroblasts overexpressing ESYT1-3xFLAG, solubilized with 1%DDM and analyzed by Blue-Native PAGE.

Consistent with these results, subcellular fractionation of mouse liver (Figure 1C) showed that endogenous ESYT1 is present in the microsomal light membrane fraction containing ER, and in the heavy membrane fraction containing mitochondria and MAMs. Further gradient-purification of the heavy membranes into MAMs and pure mitochondria revealed that ESYT1 was enriched in MAMs. Significantly, SYNJ2BP the OMM partner of ESYT1 identified by BioID, in addition to being enriched in mitochondria was also present in the MAM fraction, suggesting their potential interaction at this specific subcellular microdomain.

To further characterize the function of ESYT1, we generated a CRISPR-Cas9– mediated knock-out in human fibroblasts (KO) and fibroblasts stably overexpressing a C-terminal 3xFLAG-tagged version of ESYT1 (Figure 1D). Using these cell lines, we investigated the presence of ESYT1 complexes at MAMs. BN-PAGE analysis of DDM-solubilized heavy membrane fractions (Figure 1E) revealed that endogenous ESYT1 was present at MAMs in several large complexes, the main one at approximately 410 kDa. The specificity of these complexes was confirmed by their absence in different clones of the KO cell line. Finally, the ESYT1-FLAG overexpressing cell line demonstrated that the tagged version of ESYT1 behaved similarly to the endogenous protein, forming greater amounts of the same complexes. Together, these results show that ESYT1 and its OMM partner SYNJ2BP localize to the MAM fraction, and that ESYT1 forms high molecular weight complexes.

### ESYT1 tethers ER to mitochondria

As ESYT1 is known to tether the ER membrane to the PM (Saheki 2017) and to peroxisomes (Xiao, Luo et al. 2019), we sought to determine whether ESYT1 could similarly act as a tethering protein regulating MERCs. Using transmission electron microscopy (TEM), we analyzed the morphology and characteristics of MERCs in human control fibroblasts compared to ESYT1 KO cells and KO cells where a Myc-tagged version of ESYT1 was stably re-introduced (Figure 2A). TEM images analysis revealed that the loss of ESYT1 led to a decrease in both the number and mean length of MERCs, resulting in an overall decrease in the surface of mitochondria covered by ER membrane (Figure 1B, C). MERC morphology was completely rescued by the reintroduction of ESYT1-Myc, confirming the specificity of this phenotype. These experiments show that ESYT1 acts as a physical tether between the ER and mitochondria.

**Figure 2.**
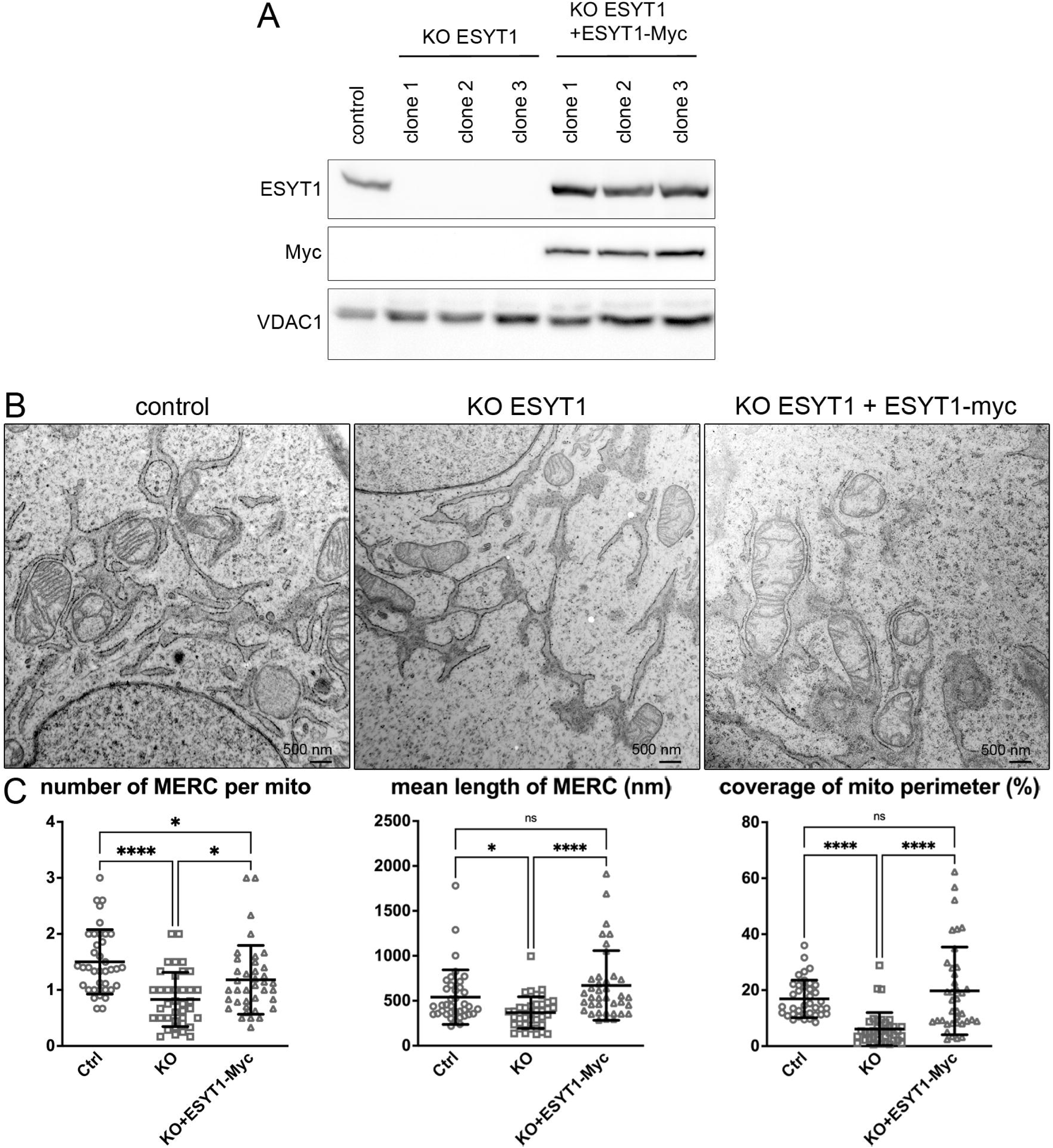
ESYT1 tethers ER to mitochondria. A. ESYT1 protein levels in control human fibroblasts, ESYT1 knock-out fibroblasts and ESYT1 knock-out fibroblasts expressing ESYT1-Myc. Whole cell lysates were analyzed by SDS-PAGE and immunoblotting. VDAC1 was used as a loading control. B. Transmission electron microscopy images of control human fibroblasts, ESYT1 knock-out fibroblasts and ESYT1 knock-out fibroblasts expressing ESYT1-Myc. C. Quantitative analysis of Mitochondria-ER contact sites (MERCs) from the TEM images: the number of MERC per mitochondria, the length of MERC (nm) and the coverage of the mitochondrial perimeter by ER (%). Results are expressed as means +/− S.D. 38 images in each condition were analyzed (n=38), totalizing 245 mitochondria for control cells, 154 mitochondria for KO cells and 224 mitochondria for rescued cells. Kruskal-Wallis and *post hoc* multiple comparisons tests were applied, ns: non-significant, * p<0.05, **** p<0.0005.

### SYNJ2BP but not ESYT1 promotes the formation of mitochondria-ER contacts

We next investigated the consequences of the overexpression of ESYT1, or its mitochondrial partner SYNJ2BP on MERC architecture. The overexpression of a 3xFLAG tagged version of ESYT1 did not influence the morphology of MERCs (Figure 3B); however, as was previously demonstrated (Nemoto and De Camilli 1999, Hung, Lam et al. 2017, Pourshafie, Masati et al. 2022), SYNJ2BP overexpression strikingly promoted the formation of MERCs (Figure 3A and B), specifically by increasing the length of individual contact between the two organelles and the mitochondrial perimeter in contact with ER in a “zipper-like” fashion (Figure 3B). Concomitantly, the ER-mitochondrial network was recruited to the perinuclear region of the cell (Figure 3A). Immunofluorescence and confocal microscopy analysis confirmed both the significant increase of MERCs and the perinuclear accumulation of the ER-mitochondrial network when SYNJ2BP was overexpressed (Figure 3C). In these conditions, we also observed that endogenous ESYT1 was recruited to MERCs, where it accumulated and formed large *foci* (Figure 3C, white arrowheads).

**Figure 3.**
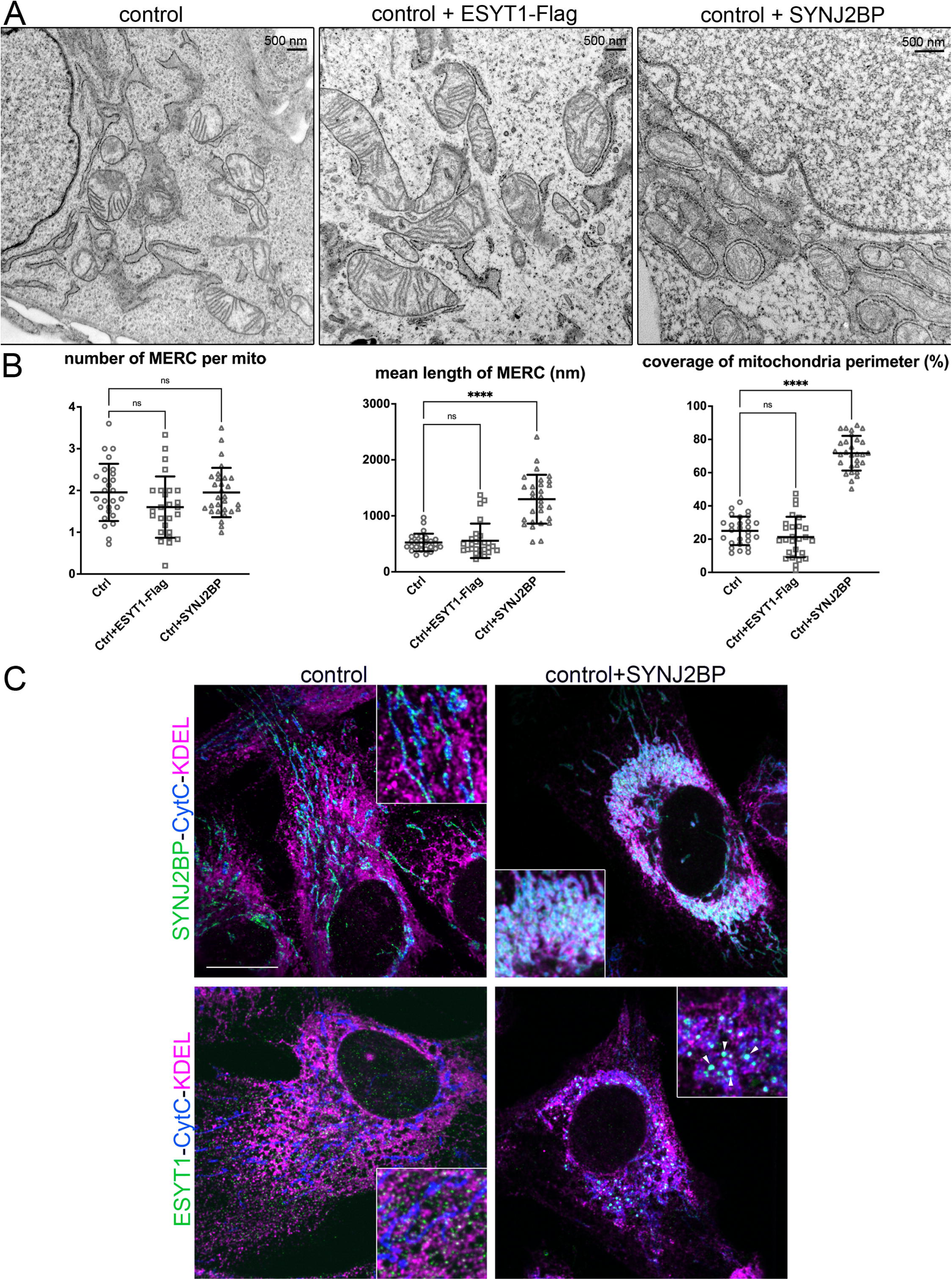
SYNJ2BP but not ESYT1 promotes the formation of mitochondria-ER contacts. A. Transmission electron microscopy images of control human fibroblasts and fibroblasts overexpressing SYNJ2BP. B. Quantitative analysis of Mitochondria-ER contact sites (MERCs) in control human fibroblasts, fibroblasts overexpressing ESYT1-Flag and fibroblasts overexpressing SYNJ2BP showing the number of MERC per mitochondria, the length of MERC (nm) and the coverage of the mitochondrial perimeter by ER (%). Results are expressed as means +/− S.D. 27 images were analyzed in control fibroblasts (n=27), totalizing 152 mitochondria. 26 images were analyzed in fibroblasts overexpressing ESYT1-Flag (n=26), totalizing 140 mitochondria. 29 images were analyzed in fibroblasts overexpressing SYNJ2BP (n=29), totalizing 300 mitochondria. Kruskal-Wallis and post hoc multiple comparisons tests were applied, ns: non-significant, **** p<0.0005. C. Confocal microscopy images of control human fibroblasts and fibroblasts overexpressing SYNJ2BP. Top panel: SYNJ2BP localization (green), cytochrome c (CytC) serves as a mitochondrial marker (cyan) and KDEL as an ER marker (magenta). Bottom panel: ESYT1 localization (green), CytC serves as a mitochondrial marker (cyan) and KDEL as an ER marker (magenta). White arrowheads highlight the accumulation of endogenous ESYT1 at MERCs when SYNJ2BP is overexpressed. Scale bar=10μm.

Due to the contribution of mitochondrial fission and fusion related proteins in the formation or stabilisation of MERCs, including MFN2 (de Brito and Scorrano 2008) and DRP1(Prudent, Zunino et al. 2015), we decided to investigate their potential contribution to the SYNJ2BP-dependent MERC formation. Control cells and cells overexpressing SYNJ2BP were depleted for either the main mitochondrial fission regulator DRP1, or the outer membrane mitochondrial fusion protein MFN2 (Figure S2). As expected, in both control cells and cells overexpressing SYNJ2BP, depletion of DRP1 led to a hyperfused mitochondrial network (Figure S2A and B), whereas loss of MFN2 induced mitochondrial fragmentation (Figure S2C and D). In both conditions, the overexpression of SYNJ2BP still promoted a strong increase of MERCs as monitored by confocal microscopy (Figure S2B and D). However, the recruitment of the ER-mitochondrial network around the nucleus was less prominent after DRP1 knockdown. We conclude that the effect of SYNJ2BP on MERC formation is independent of MFN2 and DRP1.

### SYNJ2BP is present in a high-molecular weight complex with ESYT1

To better understand the relationship between ESYT1 and SYNJ2BP, we investigated their potential interaction by BN-PAGE analysis. While endogenous SYNJ2BP ran mostly as a monomer (Figure 4A, left), when overexpressed (a condition that promotes MERCs), SYNJ2BP appeared in two high molecular weight complexes (Figure 4A, left), one of which was at the same size as the ESYT1 complex at 410 kDa (Figure 4A, right). The knock-down of SYNJ2BP slightly decreased the quantity of the ESYT1 complex, whereas its overexpression stabilized it (Figure 4A, right). A second dimension BN/SDS-PAGE analysis, confirmed that when overexpressed, a fraction of SYNJ2BP is present in two different complexes, one that runs at the size of the ESYT1 complex and one to similar size of the RRBP1 complex (Figure 4B). Knock-down of RRBP1 did not affect the assembly of ESYT1 complex (Figure 4C), nor did the knockdown of ESYT1 affect the RRBP1 complex, demonstrating that the complexes are not interdependent. However, the presence of SYNJ2BP in the 410 kDa complex is specifically dependant on ESYT1, since its depletion leads to the loss of the SYNJ2BP complex at 410 kDa (Figure 4C). These results demonstrate that ESYT1 and SYNJ2BP belong to the same complex. Huang et al. reported that the interaction of SYNJ2BP with RRBP1 depends on cytoplasmic translation activity (Hung, Lam et al. 2017). To confirm that the two SYNJ2BP complexes are independent, we analyzed the effects of puromycin, a translation inhibitor. Puromycin treatment led to a large decrease of RRBP1 protein steady state level along with an increase of ESYT1 protein level (Figure 4D). A second-dimension experiment confirmed that puromycin induced a specific loss of the SYNJ2BP-RRBP1 complex, without affecting the complex between SYNJ2BP and ESYT1 (Figure 4E). Together, these results demonstrate that SYNJ2BP interacts with both ESYT1 and RRBP1, but in two different complexes that are physically and functionally unrelated.

**Figure 4.**
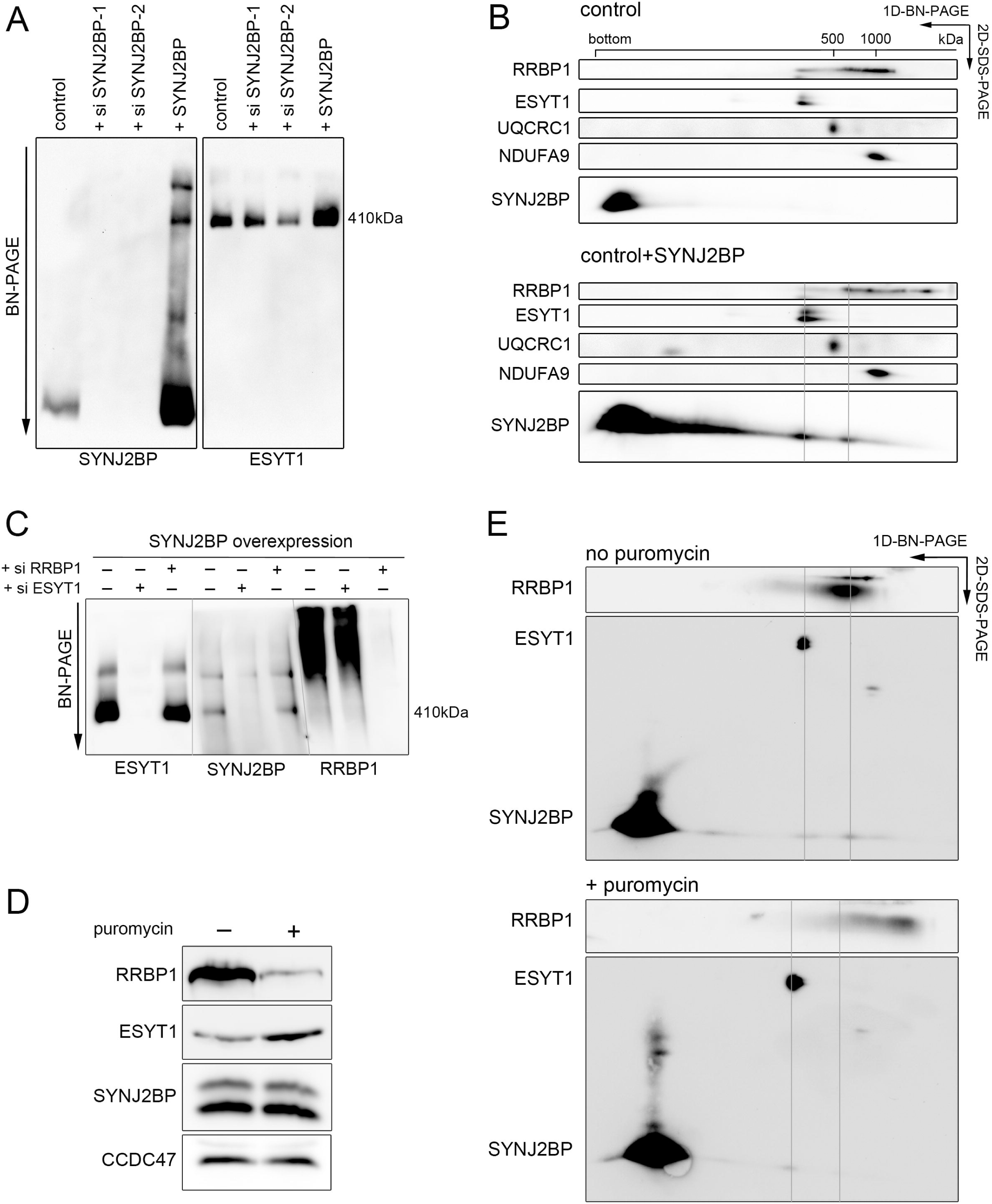
SYNJ2BP is present in a high-molecular weight complex with ESYT1. A. Characterization of ESYT1 and SYNJ2BP complexes. Heavy membrane fractions from control human fibroblasts, SYNJ2BP knocked-down fibroblasts and fibroblasts stably overexpressing SYNJ2BP were analyzed by Blue-Native PAGE. Samples were run in duplicate on the same gel and immunoblotted with anti-SYNJ2BP (left) and anti-ESYT1 antibodies (right). B. Two-dimensional electrophoresis analysis (BN-PAGE/SDS-PAGE) of SYNJ2BP interacting proteins in control human fibroblasts and fibroblasts overexpressing SYNJ2BP. The migration of known protein complexes in the first dimension is indicated on the top of the blot (UQCRC1: OXPHOS complex III at 500kDa, NDUFA9: OXPHOS complex I at 1000kDa). The position of identified SYNJ2BP containing complexes and their alignment with ESYT1 and RRBP1 containing complexes are indicated with grey lines. C. Characterization of ESYT1, SYNJ2BP and RRBP1 complexes. Heavy membrane fractions from fibroblasts overexpressing SYNJ2BP or fibroblasts overexpressing SYNJ2BP in which either ESYT1 or RRBP1 was knocked down were analyzed by Blue-Native PAGE. Samples were run in triplicate on the same gel and immunoblotted with anti-ESYT1 (left), anti-SYNJ2BP (center) and anti-RRBP1 antibodies (right). D. RRBP1, ESYT1 and SYNJ2BP protein levels in fibroblasts overexpressing SYNJ2BP untreated or treated with puromycin (200μM for 2h30). Whole cell lysates were analyzed by SDS-PAGE and immunoblotting. CCDC47 was used as a loading control. E. Two-dimensional electrophoresis analysis (BN-PAGE/SDS-PAGE) of SYNJ2BP interacting proteins in fibroblasts overexpressing SYNJ2BP untreated or treated with puromycin (200μM for 2h30). The position of identified SYNJ2BP containing complexes and their alignment with ESYT1 and RRBP1 containing complexes are indicated with grey lines.

### ESYT1 is required for ER to mitochondria Ca^2+^ transfer

In mammals, the best characterized functional feature of MERCs is Ca^2+^ flux from the ER to mitochondria required to sustain mitochondrial homeostasis. Specifically at these sites, Ca^2+^ is released from the ER through the inositol-3-phosphate receptor (IP3R) and crosses the OMM through the voltage-dependent anion channel (VDAC), which interacts with IP3R via the cytosolic protein GRP75 (Szabadkai et al., 2006). Ca^2+^ is then transported to the matrix via the inner membrane mitochondrial calcium uniporter (MCU) complex (De Stefani, Raffaello et al. 2011, Bick, Calvo et al. 2012). MERCs provide spatially constrained microdomains in which Ca^2+^ released from the ER can accumulate at concentrations sufficient to induce mitochondrial Ca^2+^ uptake via the low Ca^2+^ affinity MCU complex (Rizzuto, Pinton et al. 1998, Csordas, Renken et al. 2006, Szabadkai, Bianchi et al. 2006). As a consequence, proteins that regulate MERC formation affect ER to mitochondria Ca^2+^ transfer (de Brito and Scorrano 2008, De Vos, Morotz et al. 2012, Stoica, De Vos et al. 2014, Hirabayashi, Kwon et al. 2017).

To investigate the role of ESYT1 in mitochondrial Ca^2+^ dynamics, we compared control human fibroblasts, ESYT1 KO fibroblasts and ESYT1 KO fibroblasts expressing either ESYT1-Myc or an engineered ER-mitochondria tether reported to successfully rescue both MERCs and Ca^2+^ loss of cells devoided of the contact site protein regulators PDZD8, RMDN3-VAPB or MFN2 (Gomez-Suaga, Paillusson et al. 2017, Hirabayashi, Kwon et al. 2017, Hernandez-Alvarez, Sebastian et al. 2019). We first analyzed the capacity of mitochondria to take up exogenous Ca^2+^ upon stimulation of ER-Ca^2+^ release. Cells were transfected with the mitochondrial matrix-targeted Ca^2+^ probe reporter (mitochondrial-aequorin) (Rizzuto, Simpson et al. 1992, Granatiero, Patron et al. 2014) and mitochondrial Ca^2+^ uptake and concentrations were measured after IP3R activation with histamine to induce the release of Ca^2+^ from the ER (Figure 5). Loss of ESYT1 decreased the Ca^2+^ uptake capacities of mitochondria (Figure 5A) with a decrease of both maximal mitochondrial Ca^2+^ concentration (Figure 5B) and the rate of mitochondrial Ca^2+^ uptake (Figure 5C), all of which were rescued by re-expression of ESYT1-Myc and the engineered ER-mitochondria tether. We conclude that ESYT1-mediated MERC tethering is required for efficient ER to mitochondria Ca^2+^ transfer.

**Figure 5.**
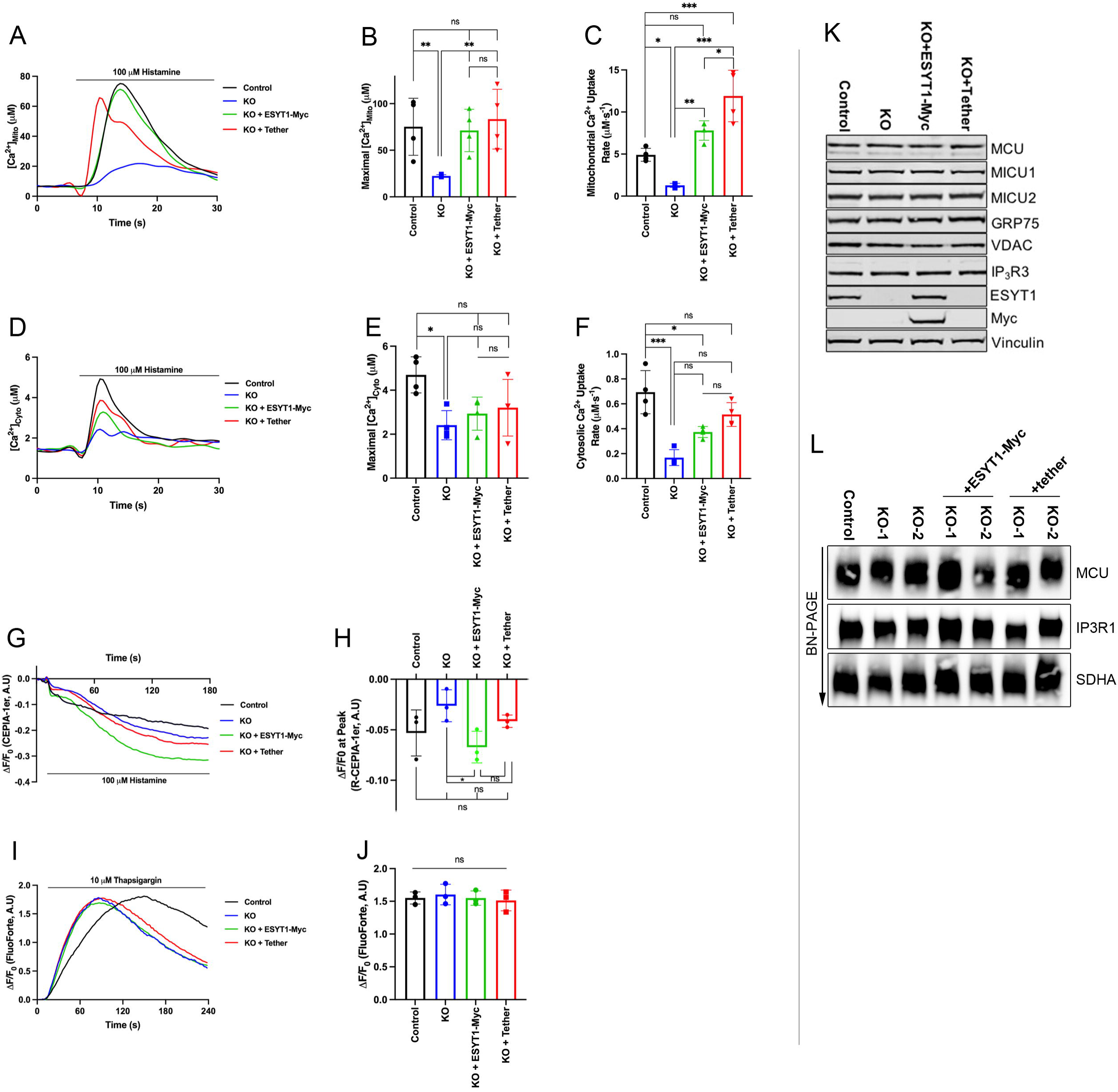
ESYT1 is required for ER to mitochondria Ca^2+^ transfer. A. Trace of mitochondrial-aequorin measurements of mitochondrial [Ca^2+^] upon histamine stimulation (100 μM) in control human fibroblasts, ESYT1 knock-out fibroblasts, ESYT1 knock-out fibroblasts expressing either ESYT1-Myc or an artificial mitochondria-ER tether. B. Quantification of maximal mitochondrial [Ca^2+^]. Results are expressed as mean ± SD. From >50 cells per condition; n=4 independent experiments. ns: not significant; **p < 0.01 (Turkey’s multiple comparisons test). C. Quantification of the rate of mitochondrial Ca^2+^ uptake. Results are expressed as mean ± SD. From >50 cells per condition; n=4 independent experiments. ns: not significant; *p < 0.05; **p < 0.01; ***p < 0.001 (Turkey’s multiple comparisons test). D. Trace of cytosolic-aequorin measurements of cytosolic [Ca^2+^] upon histamine stimulation (100 μM) in control human fibroblasts, ESYT1 knock-out fibroblasts, ESYT1 knock-out fibroblasts expressing either ESYT1-Myc or an mitochondria-ER artificial tether. E. Quantification of maximal cytosolic [Ca^2+^]. Results are expressed as mean ± SD. From >50 cells per condition; n=4 independent experiments. ns: not significant; *p < 0.05 (Turkey’s multiple comparisons test). F. Quantification of the rate of cytosolic Ca^2+^ uptake. Results are expressed as mean ± SD. From >50 cells per condition; n=4 independent experiments. ns: not significant; *p < 0.05; ***p < 0.001 (Turkey’s multiple comparisons test). G. Trace of ER [Ca^2+^] in control human fibroblasts, ESYT1 knock-out fibroblasts, ESYT1 knock-out fibroblasts expressing either ESYT1-Myc or an artificial mitochondria-ER tether. All cell lines express the ER-targeted GECI (ER-G-CEPIA1er) fluorescent probe. ER-Ca^2+^ release was stimulated with 100 μM histamine after 10 seconds of baseline (F/F0 ER-G-CEPIA1er). H. Quantification of the fold-change in fluorescence intensity (ΔF/F0) of CEPIA-1er at the initial peak induced by histamine. Results are expressed as mean ± SD; From >50 cells per condition; n=4 independent experiments. ns: not significant; *p < 0.05 (Turkey’s multiple comparisons test). I. Traces of cytosolic [Ca^2+^] in control human fibroblasts, ESYT1-KO fibroblasts and ESYT1-KO fibroblasts expressing either ESYT1-Myc or an artificial mitochondria-ER tether. All cell lines express the cytosolic fluorescent probe FluoForte. ER-Ca^2+^ release was stimulated with 10 μM thapsigargin after 10 seconds of baseline (F/F0; FluoForte). J. Quantification of the maximal fold-change in fluorescence intensity (ΔF/F0) of FluoForte upon thapsigargin stimulation (max F/F0; FluoForte). Mean ± SD, n=4 independent experiments. ns=not significant (Turkey’s multiple comparisons test). K. Whole cell lysates of control human fibroblasts, ESYT1-KO fibroblasts and ESYT1-KO fibroblasts expressing either ESYT1-Myc or an artificial mitochondria-ER tether were analyzed by SDS-PAGE and immunoblotting. Vinculin was used as a loading control. L. Heavy membrane fractions were isolated from control human fibroblasts, ESYT1 knock-out fibroblasts, ESYT1 knock-out fibroblasts expressing ESYT1-Myc or an artificial mitochondria-ER tether, solubilized and analyzed by Blue-Native PAGE. SDHA was used as a loading control.

ER-PM contact sites are responsible for store-operated Ca^2+^ entry (SOCE) (Ahmad, Narayanasamy et al. 2022). It has been suggested that ESYT1 ER-PM tethering would be activated by and reinforce SOCE (Giordano, Saheki et al. 2013, Maleth, Choi et al. 2014, Idevall-Hagren, Lu et al. 2015, Kang, Zhou et al. 2019). We therefore investigated the influence of ESYT1 loss on cytosolic Ca^2+^ concentration (Figure 5D to F) and ER-Ca^2+^ release capacity (Figure 5G and H) upon histamine stimulation. Our results showed a reduced cytosolic Ca^2+^ concentration and uptake in ESYT1-KO cells (Figure 5D-F), likely due to a decreased ER-Ca^2+^ release upon histamine stimulation (Figure 5G and H). The ER-Ca^2+^ release defect observed was fully rescued by re-expression of ESYT1-Myc but not the artificial tether, highlighting the distinct roles of ESYT1 in Ca^2+^ regulation at the ER-PM and at MERCs. Measurment of cytosolic Ca^2+^ after tharpsigargin treatment in Ca^2+^-fee media, an inhibitor of the sarco/endoplasmic reticulum Ca^2+^ ATPase SERCA that blocks Ca^2+^ pumping into the ER, showed that ESYT1 KO does not influence the total ER Ca^2+^ pool (Figure 5 I and J). Finally, immunoblot analysis (Figure 5K) showed that the levels of the major proteins involved in ER to mitochondria Ca^2+^ transfer regulation were not affected, nor was the assembly of the IP3R or the MCU complexes (Figure 5L). These results suggest that the observed ER to mitochondria Ca^2+^ transfer defects were due to decreased formation of MERCs rather than impairments in the Ca^2+^ transport machinery.

This was confirmed in HeLa cells where silencing of ESYT1 also led to a decrease of mitochondrial Ca^2+^ uptake upon histamine stimulation, monitored by genetically encoded Ca^2+^ indicator targeted to mitochondrial matrix (CEPIA-2mt) (Suzuki, Kanemaru et al. 2014). (Figure S3A and B). ESYT1 silencing in HeLa cells did not impact ER Ca^2+^ store measured by the ER-targeted R-GECO Ca^2+^ probe (Figure S3C and D) and the expression of the artificial mitochondria-ER tether was able to rescue mitochondrial Ca^2+^ defects observed in ESYT1 silenced cells (Figure S3B). Compared to the chronic loss of ESYT1, its acute silencing led to increased levels of proteins involved in ER to mitochondria Ca^2+^ transfer, including MCU, MICU2, VDAC and IP3R3 (Figure S3E). While this may reflect a compensation for the loss of ESYT1 or a response to stress, this confirms that the observed anomalies in Ca^2+^ trafficking are specifically due to MERC defects.

### SYNJ2BP is required for ER to mitochondria Ca^2+^ transfer

Based on these results, we next investigated the role of OMM ESYT1 partner SYNJ2BP in mitochondrial Ca^2+^ dynamics (Figure 6). To do so, we generated CRISPR-Cas9–mediated SYNJ2BP knock-out human fibroblasts (KO, two different clones) and fibroblasts overexpressing SYNJ2BP (either bulk cultures or a clone) and compared them to control fibroblasts (Figure 6). Similar to ESYT1 loss, the absence of SYNJ2BP strongly decreased both maximal mitochondrial Ca^2+^ concentration (Figure 6A and B) and mitochondrial Ca^2+^ uptake rate (Figure 6C). SYNJ2BP overexpression however significantly increased mitochondrial Ca^2+^ uptake capacity upon histamine stimulation (Figure 6A-C). In contrast to ESYT1, the level of SYNJ2BP did not influence cytosolic Ca^2+^ concentration (Figure 6D to F) upon histamine stimulation. However, SYNJ2BP overexpression lead to increased levels of two proteins involved in ER to mitochondria Ca^2+^ transfer MICU2 and IP3R3 (Figure 6G).

**Figure 6.**
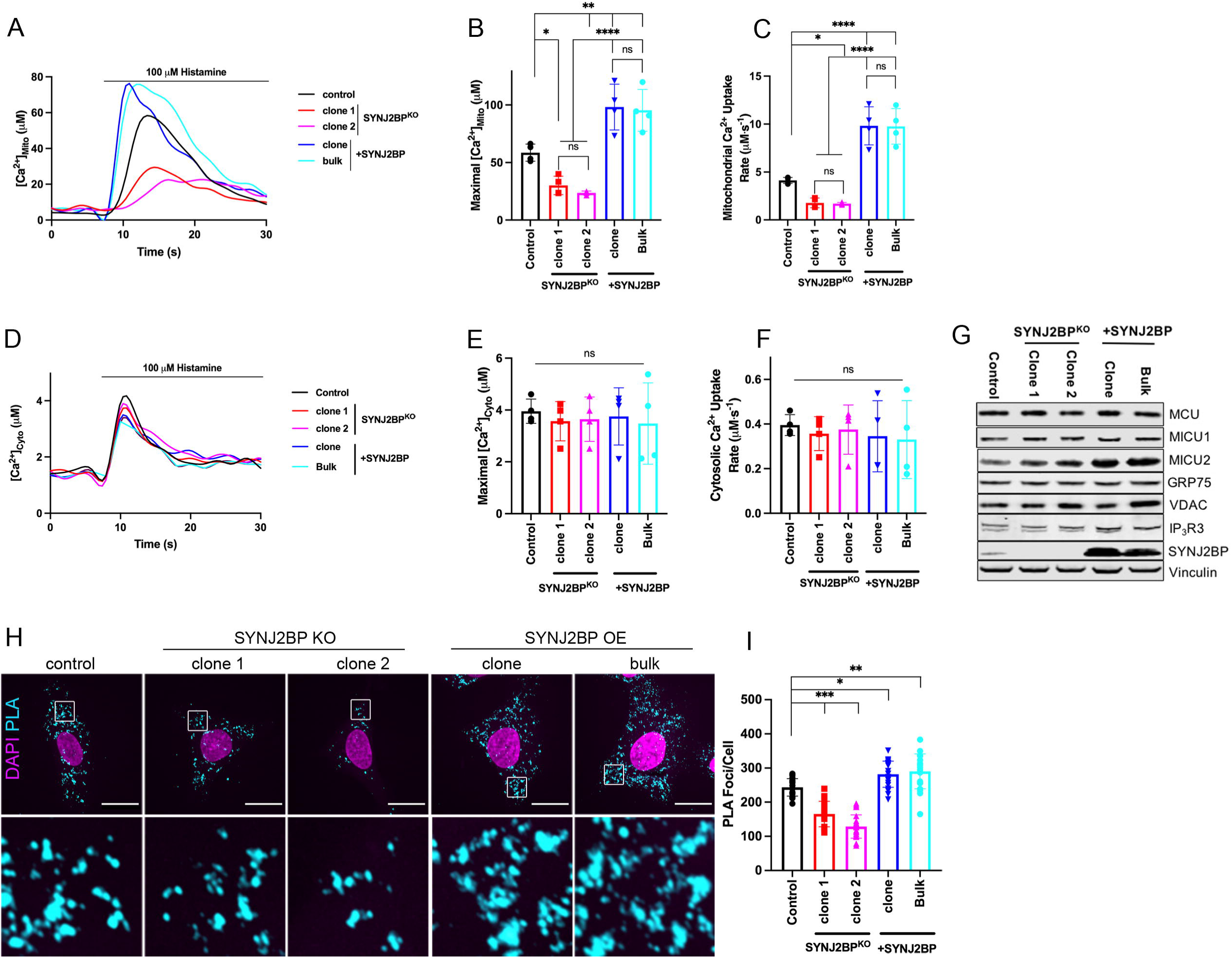
SYNJ2BP is required for ER to mitochondria Ca^2+^ transfer. A. Trace of mitochondrial-aequorin measurements of mitochondrial [Ca^2+^] upon histamine stimulation (100 μM) in control human fibroblasts, SYNJ2BP knock-out fibroblasts (clone 1 and 2) and fibroblasts overexpressing SYNJ2BP (clone and bulk). B. Quantification of maximal mitochondrial [Ca^2+^]. Results are expressed as mean ± SD. From >50 cells per condition; n=4 independent experiments. ns: not significant; *p < 0.05; **p < 0.01; ****p < 0.0001 (Turkey’s multiple comparisons test). C. Quantification of the rate of mitochondrial Ca^2+^ uptake. Results are expressed as mean ± SD. From >50 cells per condition; n=4 independent experiments. ns: not significant; *p < 0.05; ****p < 0.0001 (Turkey’s multiple comparisons test). D. Trace of cytosolic-aequorin measurements of cytosolic [Ca^2+^] upon histamine stimulation (100 μM) in control human fibroblasts, SYNJ2BP knock-out fibroblasts (clone 1 and 2) and fibroblasts overexpressing SYNJ2BP (clone and bulk). E. Quantification of maximal cytosolic [Ca^2+^]. Results are expressed as mean ± SD. From >50 cells per condition; n=4 independent experiments. ns: not significant (Turkey’s multiple comparisons test). F. Quantification of the rate of cytosolic Ca^2+^ uptake. Results are expressed as mean ± SD. From >50 cells per condition; n=4 independent experiments. ns: not significant (Turkey’s multiple comparisons test). G. Whole cell lysates of control human fibroblasts, SYNJ2BP knock-out fibroblasts (clone 1 and 2) and fibroblasts overexpressing SYNJ2BP (clone and bulk) were analyzed by SDS-PAGE and immunoblotting. Vinculin was used as a loading control. H. Representative confocal images of PLA experiment in control human fibroblasts, SYNJ2BP knock-out fibroblasts (clone 1 and 2) and fibroblasts overexpressing SYNJ2BP (clone and bulk). Anti-VDAC1 and anti-IP3R1 were used as primary antibodies in the assay. Scale bars represent 20 μm. I. Quantification of average number of PLA *foci* per cell corresponding to (H). At least 20 cells were quantified per condition per independent experiment, n=3 independent experiments. Error bars represent mean ± SD. * p < 0.05, ** p < 0.01, *** p < 0.001.

To better understand the effect of SYNJ2BP on mitochondrial Ca^2+^ uptake, we further analyzed its role in MERCs formation using an *in situ* proximity ligation assay (PLA), an established method to analyze MERCs (Figure 6H and I) (Tubbs and Rieusset 2016). As seen in our TEM analysis (Figure 3A and B), overexpression of SYNJ2BP increased the number of MERCs, monitored by the increase of the number of PLA *foci* per cell compared to controls. On the opposite, SYNJ2BP KO led to a reduction in the number of PLA *foci* per cell, indicating a decrease of MERCs (Figure 6H and I). Together these results demonstrate that the quantity of MERCs is proportional to the level of SYNJ2BP expression, which therefore influences mitochondrial Ca^2+^ uptake capacity.

### ESYT1 regulates mitochondrial lipid homeostasis

Mitochondrial lipid composition is distinct from that in other organelles (Funai, Summers et al. 2020) and plays a critical role in the regulation of mitochondrial and cellular homeostasis (Sassano, Felipe-Abrio et al. 2022, Ventura, Martinez-Ruiz et al. 2022). The most abundant mitochondrial phospholipids are phosphatidylcholine (PC), phosphatidylethanolamine (PE), cardiolipin (CL), phosphatidylinositol (PI) and phosphatidylserine (PS). CL and PE are synthetized in the IMM, requiring the import of precursors lipids, phosphatidic acid (PA) and PS, respectively, from the ER membrane at MERCs. Indeed, numerous studies have highlighted the critical contribution of MERCs in generating a platform for efficient lipid exchanges between the two organelles (Tamura, Kawano et al. 2020).

As the ESYT1-SYNJ2BP complex controls MERC architecture, we investigated the role of the SMP-domain containing protein ESYT1 in lipid transfer from ER to mitochondria. We performed shotgun mass spectrometry lipidomics, allowing broad coverage of lipids and absolute quantification (Lipotype GmbH, Dresden, Germany) from purified mitochondria. We compared control human fibroblasts (control, n=3), ESYT1 KO fibroblasts (KO, n=4) and ESYT1 KO fibroblasts expressing either ESYT1-Myc (Rescue, n=6) or the ER-mitochondria artificial tether (Tether, n=6). Over 1484 lipid entities identified and quantified of which 149 were statistically different after filtering (Table S2). Multi-variant data analysis using principal component analysis (PCA) (Figure 7A) and hierarchical clustering with heatmap analysis (Figure S4A) showed tight clustering of the replicates and a clear separation between control, KO and rescue conditions. ESYT1 and artificial tether overexpressing samples clustered together, suggesting that the mitochondrial lipid content is similar in these samples. Figure S4B shows the profile of the different lipid classes identified. The loss of ESYT1 resulted in a decrease of the three main mitochondrial lipid categories CL, PE and PI, which was accompanied by an increase in PC (Figure 7B). Importantly, re-introduction of both ESYT1 and the artificial tether rescued this phenotype. Together these results demonstrate that ESYT1 is required for lipid transfer from ER to mitochondria, likely through its tethering function as this phenotype is completely rescued by the artificial tether, suggesting that other lipid transport proteins are involved.

**Figure 7.**
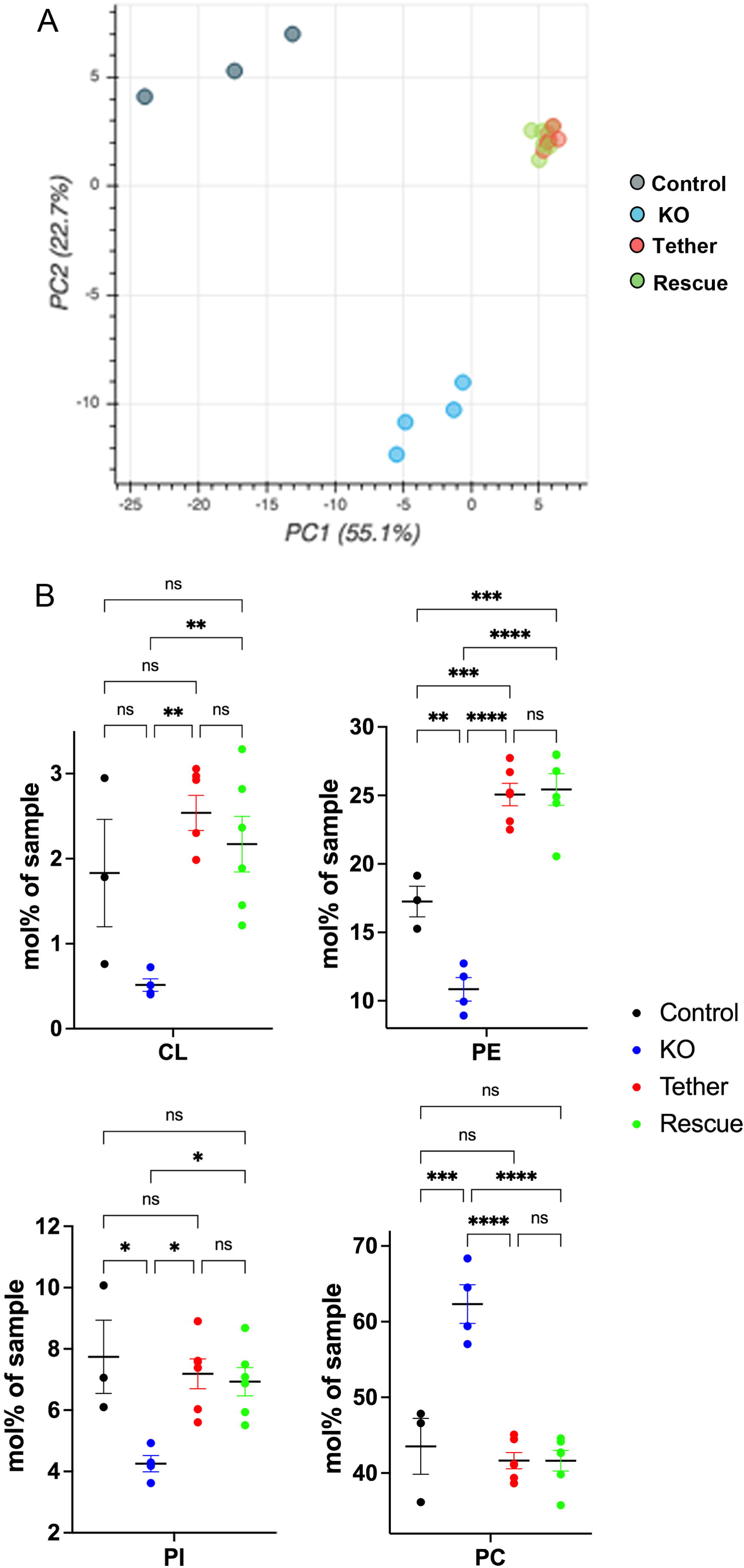
ESYT1 regulates mitochondrial lipid homeostasis. Sucrose bilayer purified mitochondria from control human fibroblasts (control, n=3), ESYT1 KO fibroblasts (KO, n=4) and ESYT1 KO fibroblasts expressing either ESYT1-Myc (Rescue, n=6) or an mitochondria-ER artificial tether (Tether, n=6) were analyzed for absolute quantification of lipid content using shotgun mass spectrometry lipidomics. A. PCA analysis of individual samples. Lipid species mol% were used as input data. B. Lipid class profile of cardiolipins (CL), phosphatidylethanolamines (PE), phosphatidylinositols (PI) and phosphatidylcholines (PC). Data are presented as molar % of the total lipid amount (mol%). One-way ANOVA with multiple comparisons analysis was applied. Error bars represent mean ± SEM. ns: not significant, * p < 0.05, ** p < 0.01, *** p < 0.001, **** p < 0.0001.

The lipid composition of the IMM influences the efficiency of oxidative phosphorylation (Funai, Summers et al. 2020). We observed that ESYT1 KO cells have a growth defect when grown on (glucose-free) galactose medium, which forces cells to rely exclusively on OXPHOS for ATP production (Figure S5). This phenotype was completely rescued by expression of Myc-tagged ESYT1 and partially rescued by expression of the artificial tether. This highlights the critical role of MERCs in tuning the mitochondrial lipid content and capacities according to cellular demand.

## DISCUSSION

This study demonstrates that the ESYT1-SYNJ2BP tethering complex regulates essential physiological functions that occur at the mitochondrial-ER interface. ESYT1 and SYNJ2BP localize to MAM subdomains where they interact in a high molecular weight complex, favouring the formation of MERCs. Loss of this tethering function results in reduced mitochondrial calcium uptake capacity, impaired mitochondrial lipid homeostasis, and slower cellular proliferation on an obligately aerobic substrate. Thus the ESYT1-SYNJ2BP complex fulfills all the essential criteria for a *bona fide* inter-organellar tether (Eisenberg-Bord, Shai et al. 2016, Scorrano, De Matteis et al. 2019). Although ESYT1 harbours calcium-binding and lipid transfer domains, both functions can be replaced by an artificial mitochondria-ER tether.

A challenge for the study of mitochondria-ER contact sites is the multiplicity of described tethers. Although one might predict that the loss of one protein complex may not be sufficient to disrupt MERC structure and function, that is in not what we observed for ESYT1-SYNJ2BP in this study. That appears to be a general observation for the other mammalian proteins that have been proposed to tether the two organelles: PDZD8 (Hirabayashi, Kwon et al. 2017), the dually OMM- and ER-localized MFN2 (de Brito and Scorrano 2008), and the OMM protein RMDN3 that interacts with the ER protein VAPB (De Vos, Morotz et al. 2012, Stoica, De Vos et al. 2014). All have been shown to regulate MERC formation and loss of function in all cases can be rescued by an engineered ER-OMM linker (Gomez-Suaga, Paillusson et al. 2017, Hirabayashi, Kwon et al. 2017, Hernandez-Alvarez, Sebastian et al. 2019) indicating that each of these protein complexes constitutes an essential tether. Whether or how the loss of one tether affects the other tethering complexes remains unexplored, but loss of individual tethers is clearly sufficient to provoke abnormal cellular calcium dynamics and interorganellar lipid transport. These data suggest, at least in the cellular models where they have been studied, that compensatory mechanisms are not commonly upregulated. This may not be the case in animal models. For instance, the loss of all three ESYTs does not affect mouse development, viability, fertility, brain structure, ER morphology, or synaptic protein composition (Sclip, Bacaj et al. 2016, Tremblay and Moss 2016), so clearly adaptive mechanisms exist. In fact, the loss of all ESYTs induces the expression the lipid transfer proteins OSBPL5 and OSBPL8 and the SOCE associated proteins ORAI1 and STIM1 (Tremblay and Moss 2016). A mechanistic resolution of the inter-relatedness of different tethering complexes will require further study.

The multiplicity of tether complexes also suggests the existence of different types of MERCs of variable composition, sustaining specific functions such as lipid transfer, calcium exchange or regulation of apoptosis. We demonstrated that contact sites occupied by SYNJ2BP and MFN2 are independent and are likely physically and functionally different since SYNJ2BP still promoted MERC formation in the absence of MFN2 (Figure S2). We also show that SYNJ2BP can be part of two different complexes, with ESYT1 or RRBP1, suggesting that it may sustain multiple functions at MERCs. Moreover, while the loss of either ESYT1 or SYNJ2BP reduces the number and length of MERCs, only the overexpression of SYNJ2BP enhanced MERC formation, leading to the recruitment of ESYT1 at MERCs (Figure 3C), stabilization of ESYT1 oligomers at MAMs (Figure 4A) and increased mitochondrial Ca^2+^ uptake capacity (Figure 6). SYNJ2BP acts like a glue zipping ER to mitochondria, the quantity of MERCs being proportional to the level of SYNJ2BP expression (Figure 6). Interestingly, it has recently been reported that SYNJ2BP dependant MERCs are involved in the physiopathology of neuronal and viral diseases (Duan, Wang et al. 2022, Pourshafie, Masati et al. 2022).

Here we show that the loss of ESYT1 altered mitochondrial lipid composition with significant decreases of CL, PE and PI that, in addition to being amongst the most abundant lipids in mitochondrial membranes (Funai, Summers et al. 2020), are essential for normal mitochondrial physiology (Belikova, Vladimirov et al. 2006, Acin-Perez, Fernandez-Silva et al. 2008, Bottinger, Horvath et al. 2012, Raemy and Martinou 2014, Hsu, Liu et al. 2015, Acoba, Senoo et al. 2020). Thus, lowering the levels of both PE and CL leads to serious growth defects of cells (Joshi, Thompson et al. 2012, Basu Ball, Neff et al. 2018), a phenotype that we observed in ESYT1 KO cells.

While ESYT1 is not actively transferring lipids from ER to mitochondria, it is essential for optimal lipid transfer through its tethering property. The mechanical tethering provided by ESYT1 might organize specialized membrane domains that serve as platforms to recruit other lipid transport proteins. Several proteins have been proposed to participate to the lipid exchange between ER and mitochondria in mammals including RMDN3 (Yeo, Park et al. 2021) and MFN2 (Hernandez-Alvarez, Sebastian et al. 2019). Of particular interest, VPS13D is present at MERCs, binds the OMM GTPase RHOT2 (Guillen-Samander, Leonzino et al. 2021) and has been proposed to link ER to mitochondria and support lipid transfer (Guillen-Samander, Leonzino et al. 2021).

OSBPL5 and OSBPL8 were shown to localize to MAMs, their loss leading to mitochondrial morphology and respiration defects (Galmes, Houcine et al. 2016). OSBPL5 and OSBPL8 actually bind to the mitochondrial intermembrane bridging (MIB)/mitochondrial contact sites and cristae junction organizing system (MICOS) complexes, where they mediate non-vesicular transport of PS from ER to mitochondria (Monteiro-Cardoso, Rochin et al. 2022). Interestingly, we found VPS13D and VPS13A as proximity interactors of SYNJ2BP. Likewise, we found OSBPL8 as a proximity interactor of ESYT1, suggesting a potential partnership between ESYT1 as a tether and the lipid transport protein OSBPL8.

The molecular mechanisms that regulate SYNJ2BP-ESYT1 complex formation remain unknown. In contrast, the function of ESYT1 at ER-PM contact sites has been extensively studied (Saheki 2017). ESYT1 consists of an N-terminal hairpin-like transmembrane domain that anchors ESYT1 to the ER. The ESYT1 SMP domain binds and transports lipids *in vitro* (Bian, Saheki et al. 2018) and the five C2 domains (A to E) bind Ca^2+^ and mediate interactions with phospholipids (Corbalan-Garcia and Gomez-Fernandez 2014). Ca^2+^ binding to the C2C domain in ESYT1 enables the binding of the C2E domain to PI(4,5)P_2_-rich membranes, either at the PM or at peroxisomes. Whether similar PI(4,5)P_2_-rich domains exist at the surface of mitochondria regulating ESYT1 recruitment remains an open question.

SYNJ2BP is a C-terminal tail-anchored OMM protein with a PDZ domain facing the cytosol (Hung, Lam et al. 2017). PDZ domains are small globular protein–protein interaction domains that bind the C-terminus of partner proteins. Some PDZ domains can also bind phosphatidylinositides, especially PI(4,5)P_2_ and cholesterol (Liu and Fuentes 2019), suggesting a synergistic binding of PDZ to phosphatidylinositide lipids and proteins (Pemberton and Balla 2019). This raises the possibility that the binding of ESYT1 to SYNJ2BP could involve an interaction with PI(4,5)P_2_ at the surface of the

OMM, an hypothesis that will require further investigation.

## Supporting information

BioID analysis

Lipidomics analysis

## ACKNOWLEDGMENTS

We thank Kathleen Daigneault, Isabella Straub and Hanish Anand (MRC MBU) for advice and excellent technical assistance.

## AUTHOR CONTRIBUTIONS

A.J. led the project, designed and performed experiments and co-wrote the manuscript. J.L.M. performed calcium measurement experiments. M.K. contributed to TEM images analysis. H.A. contributed to BioID data analysis and co-wrote the manuscript. M.J.A. performed biochemical analysis experiments. Z.Y.L. performed the BioID experiments. A.C.G. designed the BioID experiments. J.P. designed calcium measurement experiments. E.A.S. helped with experimental design, supervised the project and cowrote the manuscript.

## STAR METHODS

### Cell culture

Fibroblasts, HeLa cells, Flp-In T-REx 293 (Invitrogen) and Phoenix packaging (a kind gift of Garry P Nolan) cell lines were grown in 4.5g/l glucose DMEM (Wisent 319-027-CL) supplemented with 10% fetal bovine serum in 5% CO_2_ incubator at 37°C. Galactose media was composed of DMEM (Gibco, A14430-01) supplemented with 10% dialysed fetal bovine serum, sodium pyruvate (Sigma), MEM non-essential amino acids (Gibco), GlutaMAX (Gibco) and 4.5g/l of galactose. Cell lines were regularly tested for mycoplasma contamination. For cytosolic translation inhibition, cells were treated with puromycin at 200 μM final concentration for 2.5 hours. ON-TARGETPlus SMARTPool siRNA (Dharmacon) were used for transient knockdown of *DRP1* (L-012092-00-0005) and *MFN2* (L-012961-00-0005) and stealth siRNA (Invitrogen) for knockdown of *SYNJ2BP* (HSS124399), *RRBP1* (HSS109381) and *ESYT1* (HSS146329). siRNAs were transiently transfected into cells using Lipofectamine RNAiMAX (Invitrogen), according to the manufacturer’s specifications. Cells were analyzed after 6 days.

### Generation of knock-out and overexpression cell lines

Knock-out cell lines of ESYT1 and SYNJ2BP were generated by CRISPR-Cas9–mediated gene editing in human fibroblast cells. Gene-specific target sequence 5’GTTCTTTCTCGTCGCGGACC3’ for *ESYT1* and 5’GAAGAGATCAATCTTACCAG3’ for *SYNJ2BP* was cloned into pSpCas9(BB)-2A-Puro (PX459) V2.0 (62988; Addgene) (Ran, Hsu et al. 2013) and transfected into cells by Lipofectamine 3000 (Thermo Fisher Scientific) according to the manufacturer’s instructions. The day after, transfected cells were selected by addition of puromycin (2.5 μg/ml) for 2 days. Individual clones were screened for loss of target protein by immunoblotting and frameshift mutations were confirmed by genomic sequencing. Cells stably overexpressing ESYT1-3xFLAG, ESYT1-Myc, SYNJ2BP and the artificial tether (blue fluorescent protein (BFP) with OMM targeting sequence of mAKAP1 at the N terminus and the ER targeting sequence of yUBC6 at the C terminus (Hirabayashi, Kwon et al. 2017)) were engineered by retroviral infection of virus produced in Phoenix cells transfected with pLXSH-Hygro plasmids as described previously (Weraarpachai, Antonicka et al. 2009). Flp-In T-REx 293 stable cell lines were generated as previously described (Antonicka, Lin et al. 2020).

### Bait cloning

All constructs were generated using Gateway cloning into a suitable pDEST-pcDNA5-BirA*-FLAG construct (to create either an N- or C-terminal BirA*-FLAG fusion proteins). Gateway entry clones for *ESYT1* (GeneCopoeia, cat. # HOC21918), *ESYT2* (Addgene, #66831), *PDZD8* (DNasu, HsCD00400023) and *TEX2* (DNasu, HsCD00351688) were used. For *SYNJ2BP*, an entry clone was created by PCR amplification of the ORF from human cDNA (fwd primer: 5’- GGGGACAAGTTTGTACAAAAAAGCAGGCTTCATGAACGGAAGAGTGGATTATTTG- 3’, rev primer: 5’- GGGGACCACTTTGTACAAGAAAGCTGGGTTCAAAGTTGTTGCCGGTATCT-3’), followed by a sub-cloning into pDONR-221 (Invitrogen).

For selection of stable Flp-In T-REx 293 expressing clones a previously described procedure was used, and representative images for all baits are shown in Figure S1 (Antonicka, Lin et al. 2020).

### Immunofluorescence

For immunofluorescence experiments, cells plated on coverslips 24 hours prior to experiment were fixed using 4% formaldehyde in PBS for 20 min at 37°C. Coverslips were washed 3 times with PBS and cells were permeabilized in 0.1% Triton in PBS for 15 min at room temperature. Following three washes with PBS, coverslips were blocked in PBS containing 5% BSA for 30 minutes, incubated with primary antibodies for one hour at room temperature, washed 3 times with PBS and incubated with Alexa conjugated secondary antibodies (1:2000) and DAPI (1:2000) for 30 min at room temperature. Coverslips were washed three times with PBS and mounted with Fluromount-G (Thermofisher). Cells were imaged with Olympus IX83 microscope connected with Yokogawa CSU-X confocal scanning unit, using UPLANSAPO 100x/1.40 Oil objective (Olympus) and Andor Neo sCMOS camera. Images were processed in Fiji (Schindelin, Arganda-Carreras et al. 2012).

### BioID sample preparation, mass-spec data acquisition and MS data analysis

BioID analysis, mass spectra acquisition and MS data analysis was performed as described previously (Antonicka, Lin et al. 2020). For analysis with SAINT, only proteins with iProphet protein probability >0.95 were considered, which corresponds to an estimated protein level FDR of ~0.5%. A minimum of two detected peptide ions was required. SAINTexpress analysis was performed using version exp3.6.3 with two biological replicates per bait. SAINT analysis included 48 negative control runs used previously in (Antonicka, Lin et al. 2020) consisting of untransfected Flp-In T-Rex 293 cells (to detect endogenously biotinylated proteins) and BirA*-FLAG-GFP cells (to detect preys that become promiscuously biotinylated). A threshold at 1% Bayesian false discovery rate (BFDR) was used to select high-confidence proximity interactors (Table S1). All non-human protein contaminants were removed from the SAINT file.

### Databases used for analysis

Mitocarta 3.0 (Rath, Sharma et al. 2021) was used for annotation of detected preys as mitochondrial proteins. PANTHER17.0 database was used for Gene Ontology annotations (GO database released 22/03/2022).

### BioID data visualization

A file combining BioID data of all SMP-domain proteins was used as input file for ProHits-viz (Knight, Choi et al. 2017) analysis and the specificity module was used with average spectra (AvgSpec) as the abundance measure. Subtraction of the spectral counts across the controls was performed. The spectral counts for each prey were normalized to the Prey Sequence Length. The figures were annotated and color-coded using the visualization module of ProHits-viz. Venn diagrams were created using Venny 2.1 (https://bioinfogp.cnb.csic.es/tools/venny/index.html).

### Mouse liver fractionation

C57/Bl6N male mice were obtained from Jackson Laboratories, and liver harvesting and animal handling were approved and performed in accordance with the Montreal Neurological Institute Animal Care Committee regulations. The fractionation was performed as described in (Aaltonen, Alecu et al. 2022).

### Heavy-membrane preparation and sucrose bilayer mitochondrial purification

For heavy membrane fraction preparation, cells were rinsed twice, resuspended in icecold ST buffer (250 mM sucrose, 10 mM Tris–HCl pH 7.4) + Complete protease inhibitor cocktail (Roche) and homogenized with ten passes of a prechilled, zero-clearance homogenizer (Kimble/Kontes). A post-nuclear supernatant was obtained by centrifugation of the samples twice for 10 min at 600 × *g*. Heavy membranes were pelleted by centrifugation for 10 min at 10,000 × *g* and washed once in the same buffer. Protein concentration was determined by Bradford assay.

For sucrose bilayer mitochondrial purification, heavy membranes fractions were resuspended in ST buffer, loaded on top of a sucrose bilayer (1ml of 1M sucrose in ST buffer on top of 1ml of 1.7M sucrose in ST buffer) and centrifuged for 40 min at 70.000g. The band at the sucrose bilayer intersection containing pure mitochondria was harvested, diluted in ST buffer and centrifuged for 10min at 12.000g. The pellet was then washed once with ST buffer. Protein concentration was determined by Bradford assay.

### SDS-PAGE, BN-PAGE, two-dimensional electrophoresis and Western blot

Blue-Native PAGE (BN-PAGE) was used to separate individual protein complexes. Heavy membranes were solubilized with 1% dodecyl maltoside or 8 mg/ml of digitonin for MCU and IP3R complexes. Solubilized samples (10–20 μg) were run in the first dimension on 6%–15% polyacrylamide gradient gels as described in detail previously (Leary and Sasarman 2009). For the second-dimension analysis, BN-PAGE/SDS-PAGE was carried out as detailed previously (Antonicka, Ogilvie et al. 2003).

SDS-PAGE was used to separate denatured whole-cell extracts, heavy membranes or mouse fractionation samples. In general, whole cells were extracted with 1.5% lauryl maltoside in PBS, after which 20 μg of protein was run on either 10%, 12%, or 15% polyacrylamide gels.

Separated proteins were transferred to a nitrocellulose membrane (PALL), and subsequently incubated with indicated primary and secondary antibodies in 5% skim-milk Tris-buffered saline solution with 0.1% Tween 20.

### TEM analysis

Cells were washed in 0.1 M Na cacodylate washing buffer (Electron Microscopy Sciences) and fixed in 2.5% glutaraldehyde (Electron Microscopy Sciences) in 0.1 M Na cacodylate buffer overnight at 4°C. Cells were then washed three times in 0.1 M Na cacodylate washing buffer for a total of 1 h, incubated in 1% osmium tetroxide (Mecalab) for 1 h at 4°C, and washed with ddH_2_O three times for 10 min. Then, dehydration was performed in a graded series of ethanol/deionized water solutions from 30 to 90% for 8 min each, and 100% twice for 10 min each. The cells were then infiltrated with a 1:1 and 3:1 Epon 812 (Mecalab):ethanol mixture, each for 30 min, followed by 100% Epon 812 for 1 h. Cells were embedded in the culture wells with 100% Epon 812 and polymerized overnight in an oven at 60°C. Polymerized blocks were trimmed and 100 nm ultrathin sections were cut with an Ultracut E ultramicrotome (Reichert Jung) and transferred onto 200-mesh Cu grids (Electron Microscopy Sciences). Sections were post-stained for 8 min with 4% aqueous uranyl acetate (Electron Microscopy Sciences) and 5 min with Reynold’s lead citrate (Fisher Scientific). Samples were imaged with a FEI Tecnai-12 transmission electron microscope (FEI Company) operating at an accelerating voltage of 120 kV equipped with an XR-80C AMT, 8 megapixel CCD camera. Based on the images, MERCs characteristics (number, length, mitochondrial perimeter coverage) were measured using ImageJ software. The distance between ER and outer mitochondrial membrane (OMM) was selected within 10-80□nm, manually traced and quantified using ImageJ software.

### Proximity ligation assay

A proximity ligation assay (PLA) (Duolink Proximity Ligation Assay, Merk) was used to analyze the interaction of characterised ER and mitochondria resident proteins, which interact at MAMs, namely VDAC1 (Abcam, ab14734) and IP3R1 (Abcam, ab264281) (Tubbs and Rieusset 2016). Cells were cultured on coverslips in 24-well plates and were fixed in 5 % PFA for 10 min at 37°C, quenched using 50 mM ammonium chloride and permeabilized with 0.1 % Trition-X100 in PBS for 10 minutes. Between each step, cells were washed three times in PBS. Cells were blocked in Duolink blocking solution and incubated in a humidified chamber at 37°C for 1 hr. Primary antibodies were diluted in Duolink antibody diluent and incubated at 4°C overnight. The next day, cells were washed twice with PBS for 5 min and probed with the appropriate secondary antibodies coupled to the template DNA strands at 37°C for 1 hr at RT. The template DNA strand on each antibody was ligated by a DNA ligase at 37°C for 30 min at RT. Cells were washed twice with PBS for 5 min at RT and rolling loop DNA amplification was then initiated using a DNA polymerase and fluorescent nucleotides enabling detection by confocal microscopy. Cells were washed twice in PBS for 10 min and once in ddH_2_O for 1 min before being mounted onto glass slides using mounting media containing 4’, 6- diamidino-2- phenylindole (DAPI) (ProLong Diamond, Invitrogen). At least 20 cells were analyzed from three independent experiments.

### Lipid extraction for mass spectrometry lipidomics

Mass spectrometry-based lipid analysis was performed by Lipotype GmbH (Dresden, Germany) as described (Sampaio, Gerl et al. 2011). Lipids were extracted using a two-step chloroform/methanol procedure (Ejsing, Sampaio et al. 2009). Samples were spiked with internal lipid standard mixture containing: cardiolipin 16:1/15:0/15:0/15:0 (CL), ceramide 18:1;2/17:0 (Cer), diacylglycerol 17:0/17:0 (DAG), hexosylceramide 18:1;2/12:0 (HexCer), lyso-phosphatidate 17:0 (LPA), lyso-phosphatidylcholine 12:0 (LPC), lyso-phosphatidylethanolamine 17:1 (LPE), lyso-phosphatidylglycerol 17:1 (LPG), lyso-phosphatidylinositol 17:1 (LPI), lyso-phosphatidylserine 17:1 (LPS), phosphatidate 17:0/17:0 (PA), phosphatidylcholine 17:0/17:0 (PC), phosphatidylethanolamine 17:0/17:0 (PE), phosphatidylglycerol 17:0/17:0 (PG), phosphatidylinositol 16:0/16:0 (PI), phosphatidylserine 17:0/17:0 (PS), cholesterol ester 20:0 (CE), sphingomyelin 18:1;2/12:0;0 (SM), triacylglycerol 17:0/17:0/17:0 (TAG). After extraction, the organic phase was transferred to an infusion plate and dried in a speed vacuum concentrator. 1st step dry extract was re-suspended in 7.5 mM ammonium acetate in chloroform/methanol/propanol (1:2:4, V:V:V) and 2nd step dry extract in 33% ethanol solution of methylamine in chloroform/methanol (0.003:5:1; V:V:V). All liquid handling steps were performed using Hamilton Robotics STARlet robotic platform with the Anti Droplet Control feature for organic solvents pipetting.

### Lipidomics MS data acquisition

Samples were analyzed by direct infusion on a QExactive mass spectrometer (Thermo Scientific) equipped with a TriVersa NanoMate ion source (Advion Biosciences). Samples were analyzed in both positive and negative ion modes with a resolution of Rm/z=200=280000 for MS and Rm/z=200=17500 for MSMS experiments, in a single acquisition. MSMS was triggered by an inclusion list encompassing corresponding MS mass ranges scanned in 1 Da increments (Surma, Herzog et al. 2015). Both MS and MSMS data were combined to monitor CE, DAG and TAG ions as ammonium adducts; PC, PC O-, as acetate adducts; and CL, PA, PE, PE O-, PG, PI and PS as deprotonated anions. MS only was used to monitor LPA, LPE, LPE O-, LPI and LPS as deprotonated anions; Cer, HexCer, SM, LPC and LPC O- as acetate adducts.

### Lipidomics data analysis and post-processing

Data were analyzed with Lipotype’s in-house developed lipid identification software based on LipidXplorer (Herzog, Schwudke et al. 2011, Herzog, Schuhmann et al. 2012). Data post-processing and normalization were performed using Lipotype’s in-house developed data management system. Only lipid identifications with a signal-to-noise ratio >5, and a signal intensity 5-fold higher than in corresponding blank samples were considered for further data analysis.

### Lipidomics statistical analysis

Lipidomics result analysis was performed using the integrative tool LipotypeZoom from Lipotype. Lipids were selected with a cut-off of fold change 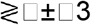 and a P-value <0.05 with a Benjamini & Hochberg adjustment.

### Aequorin-based mitochondrial and cytosolic calcium measurements

To measure cytosolic or mitochondrial Ca^2+^ concentration, cells were cultured in white 96-well plates (Corning) and reverse-transduced with adenovirus containing either the mutated mitochondrial matrix-targeted (mtAEQmut) (Montero et al., 2000) or wild-type cytosolic aequorin (CytAEQ) (Brini et al., 1995) probes and incubated overnight at 37°C and 5 % CO_2_. Cells were washed three times in balanced salt solution (BSS) + Ca2+ (120 mM NaCl, 5.4 mM KCl, 0,8 mM MgCl_2_, 6 mM NaHCO_3_, 5.6 mM D-glucose, 2 mM CaCl_2_, and 25 mM HEPES [pH 7.3]) and incubated with 5 μM coelenterazine (Sigma) in BSS + Ca^2+^ for 90 min at 37°C and 5 % CO_2_. Post-incubation, cells were washed once in BSS + Ca^2+^ and luminescence was measured by spectrophotometry (ClarioSTAR, BMG LabTek). Luminescence was measured every 2 s for 2 min. Basal luminescence was measured for 10 s followed by 100 μM histamine stimulation. At 1 min, cells were digitonized and saturated with Ca^2+^ by injection of 100 μM digitonin and 10 mM CaCl_2_, to discharge all luminous potential. Aequorin luminescence was calibrated into Ca^2+^ concentration using *equation 1*. For mtAEQmut: n = 1.43, K_TR_ = 22008 & K_R_ =22770000. For CytAEQ: n = 2.99, K_TR_ = 120 & K_R_ =7230000. Statistical significance was determined from four independent experiments (N=4) by repeated measures one-way ANOVA and Tukey’s *post hoc* test for differences.

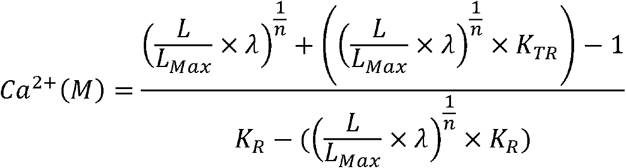

*Equation 1. Relationship between Ca^2+^ concentration and AEQ luminescence*.

*L=Light intensity, L_Max_=Sum of all light intensities, K_R_= Constant for Ca^2+^-bound state, K_TR_=Constant for Ca^2+^-unbound state, λ=Rate constant for AEQ consumption at Ca^2+^saturation. n=Number of Ca^2+^ binding sites*. (Bonora *et al*., 2013).

### Intracellular calcium analysis

Cells were seeded on a Nunc Lab-Tek chambered 8-well cover glass (Thermo Scientific). To measure mitochondrial, cytosolic and ER calcium content, cells were transfected respectively with plasmids encoding mitochondria-targeted GECI (CEPIA2mt), cytosolic-targeted GECI (R-GECO) or cytosolic-targeted FluoForte and ER-targeted GECI (R-CEPIA1er) (Suzuki, Kanemaru et al. 2014) using Fugene HD, following manufacturer’s instructions. 24 hours following transfections, cells were washed three times in a balanced salt solution buffer (BSS) (120 mM NaCl, 5.4 mM KCl, 0.8 mM MgCl_2_, 6 mM NaHCO_3_, 5.6 mM D-glucose, 2 mM CaCl_2_, and 25 mM HEPES [pH 7.3]) before analysis. Fluorescence values were then collected every 2s, and cells were stimulated with 10 μM histamine in BSS. Fluorescence was recorded for 3 minutes using the 40x objective of the Nikon Eclipse Ti-E microscope of the Andor Dragonfly spinning disk confocal system coupled with an Andor Ixon camera, exciting with 488 nm or 568 nm laser for CEPIA-2mt/G-CEPIA1ER or R-GECO respectively. Changes of fluorescence (ΔF) from each fluorescent calcium probe were normalized by basal signals before histamine stimulation (F0).

### Proliferation Assay

10,000 cells were seeded in a six wells culture dish at day 0. After 2 and 4 days of culture, cells were trypsinized, homogenized, and counted using a Neubauer counting chamber.

