## Supplemental figure legends for "ESYT1 tethers the endoplasmic reticulum to mitochondria and is required for mitochondrial lipid and calcium homeostasis"

**SUPPLEMENTAL INFORMATION**

**Table S1. BioID proximity interactome data for selected SMP-domain containing proteins and SYNJ2BP.**

Tabs A-G: Bait name indicated by the gene name_the orientation of BirA*-FLAG tag (N or C). Bait indicates the bait gene name, isoform and the orientation of BirA*-FLAG tag (N or C; column A); Prey (column B) is the NCBI protein accession number, PreyGene (column C) is the Symbol as per NCBI Gene; Mitocarta3 (column D) identifies a presence ("1") or absence ("0") in the Mitocarta 3.0 list of mitochondrial proteins; Spectral counts for each prey (column E, separated by "I" delimiter; column F, the sum of spectral counts; column G, the average of spectral counts), number of replicates performed (column H), spectral counts for the prey across all negative controls (column I), Averaged probability across replicates (AvgP; column J), maximal probability (column K), SAINT score (probability of protein-protein interaction, column L), Fold Change (counts in the purification divided by counts in the controls plus a small factor to prevent division by 0; column M), Bayesian FDR (column N), Prey Sequence Length (column O) are listed for each bait-prey relationship and are directly from the SAINTexpress output.

Note that the following entries were also manually removed from the final high confidence list associated with this manuscript: 1) "DECOY" sequences (removed during SAINTexpress) and 2) all non-human protein contaminants

Tab H: Preys identified as proximity interactors of all SMP baits (related to Figure S1). PreyGene (column A) is the Symbol as per NCBI Gene; Localization (column B) was determined based on PANTHER and GO term Cellular Compartment analysis.

Tab I: GO term functional analysis: Cellular Compartment term analysis SYNJ2BP BioID data. Source (column A) is the GeneOntology database source: Cellular Compartment (GO:CC); Term_name indicates the Cellular Compartment term (column B); Term_id (column C) is the GO term ID; Adjusted_p_value (column D) indicates the significance value; Term_size (column E) indicates how many genes are present in the GO term family; Query_size (column F) is the size of the query list; Intersection_size (column G) indicates how many query genes are present in the term list; Genes (column H) is the list of genes in the Intersection.

**Table S2. Lipidomics analysis.**

The lipidomic analysis was done by the Lipotype Shotgun Lipidomics platform (Dresden, Germany). The samples were extracted and analyzed through high resolution Orbitrap mass spectrometry and quantified and identified by Lipotype’s in house software LipotypeXplorer. All values are indicated as 'mol % of sample'. The samples are indicated as control human fibroblasts (Control_1, Control_2, Control_3, n=3), ESYT1 knock-out fibroblasts (KO_1, KO_2, KO_3, KO_4, n=4), ESYT1 knock-out fibroblasts expressing ESYT1-Myc (Rescue_1, Rescue_2, Rescue_3, Rescue_4, Rescue_5, Rescue_6, n=6), ESYT1 knock-out fibroblasts expressing a mitochondria-ER artificial tether (Tether_1, Tether_2, Tether_3, Tether_4, Tether_5, Tether_6, n=6)

Tab Species not filtered: 1484 identified lipids when no filtering and no testing for significance was applied.

Tab Species: 149 lipid species were significantly different (P-value<0.05%) with a fold change of >3. A heatmap is shown in Figure S4A.

Tab Classes: 149 lipid species were significantly different (P-value<0.05%) with a fold change of >3. The classes of those lipids are summarized and shown as a graph in Figure S4B.

**Figure S1. BioID analysis.**

A. Representative images of immunofluorescence analysis of individual baits. FLAG staining representing the bait is in green, mitochondrial marker TOMM20 in magenta, DAPI in blue. Scale bar, 10 µm.

B. Venn diagram of the BioID results for all four SMP-domain proteins.

C. Specificity plot of ESYT1-N-ter indicates the specific proximity interaction with SYNJ2BP. The specificity signifies the fold enrichment of the spectral counts detected for each prey in ESYT1 BioID compared to the spectral counts for that prey with all other baits in the dataset. Prey names for the most specific preys and for preys with the highest length-normalized spectral counts are indicated. Preys are colour-coded based on their GO term cellular compartment analysis. Mitocarta3.0 proteins are SYNJ2BP, FKBP8 and ALDH3A2.

**Figure S2. SYNJ2BP effect on MERCs is independent of the mitochondrial fission-fusion machinery.**

Confocal microscopy images of human fibroblasts. CytC serves as a mitochondrial marker (green), HSPA5 as an ER marker (magenta) and nuclei are stained with DAPI (blue). Knock-down of DRP1 in control fibroblasts (A) and in SYNJ2BP overexpressing fibroblasts (B). Knock-down of MFN2 in control fibroblasts (C) and in fibroblasts overexpressing SYNJ2BP (D). Scale bar=10μm.

**Figure S3. Silencing of ESYT1 impairs ER to mitochondria Ca^2+^ flux.**

A. Trace of mitochondrial [Ca^2+^] upon histamine stimulation (100 μM) in control HeLa cells, cells knocked-down for ESYT1 and cells knocked-down for ESYT1 that express an artificial ER-mitochondria tether. All cells express the mitochondrial Ca^2+^ probe, CEPIA-2mt.

B. Quantification of the maximal fluorescence intensity fold-change (ΔF/F0) of CEPIA-2mt induced by histamine. Results are expressed as mean ± SD; From >50 cells per condition; n=3 independent experiments. ns: not significant; *p < 0.05 (Turkey’s multiple comparisons test).

C. Trace of cytosolic [Ca^2+^] upon thapsigargin treatment (10 μM) in control HeLa cells, cells knocked-down for ESYT1 and cells knocked-down for ESYT1 that express an artificial ER-mitochondria tether. All cells express the cytosolic Ca^2+^ probe, R-GECO.

D. Quantification of the maximal fluorescence intensity fold-change (ΔF/F0) of R-GECO upon thapsigargin treatment. Results are expressed as mean ± SD; From >50 cells per condition; n=3 independent experiments. ns: not significant (Turkey’s multiple comparisons test).

E. Whole cell lysates of control HeLa cells, cells knocked-down for ESYT1 and cells knocked-down for ESYT1 that express an artificial ER-mitochondria tether were analyzed by SDS–PAGE and immunoblotting. Vinculin was used as a loading control.

**Figure S4. Mitochondrial lipids analysis.**

Sucrose bilayer purified mitochondria from control human fibroblasts (control, n=3), ESYT1 KO fibroblasts (KO, n=4) and ESYT1 KO fibroblasts expressing either ESYT1-Myc (Rescue, n=6) or an ER-mitochondria artificial tether (Tether, n=6) were analyzed for absolute quantification of lipid content using shotgun mass spectrometry lipidomics.

A. Hierarchical clustering with heatmap analysis of samples (rows) and lipids (columns).

B. Lipid class profile of analyzed samples. Data are presented as molar % of the total lipid amount (mol%). TAG: triacylglycerol; DAG: diacylglycerol; CL: cardiolipin; PA: phosphatidate; PC: phosphatidylcholine; PC-O: phosphatidylcholine ether; PE: phosphatidylethanolamine; PE-O: phosphatidylethanolamine ether; PG:

phosphatidylglycerol; PI: phosphatidylinositol; PS: phosphatidylserine; LPC: lyso-phosphatidylcholine; LPE: lyso-phosphatidylethanolamine; SM: sphingomyelin; HexCer: hexosylceramide.

**Figure S5. Cell growth curve in (glucose-free) galactose medium.**

Control human fibroblasts, ESYT1 KO fibroblasts and ESYT1 KO fibroblasts expressing either ESYT1-Myc or an ER-mitochondria artificial tether were grown in glucose-free galactose medium. 10 000 cells were plated at day 0 and grown for 4 days. n=3, 2-way Anova multiple comparisons tests were applied, * p<0.05.
