## Supplementary figures and images for "ESYT1 tethers the endoplasmic reticulum to mitochondria and is required for mitochondrial lipid and calcium homeostasis"

### Figure S1

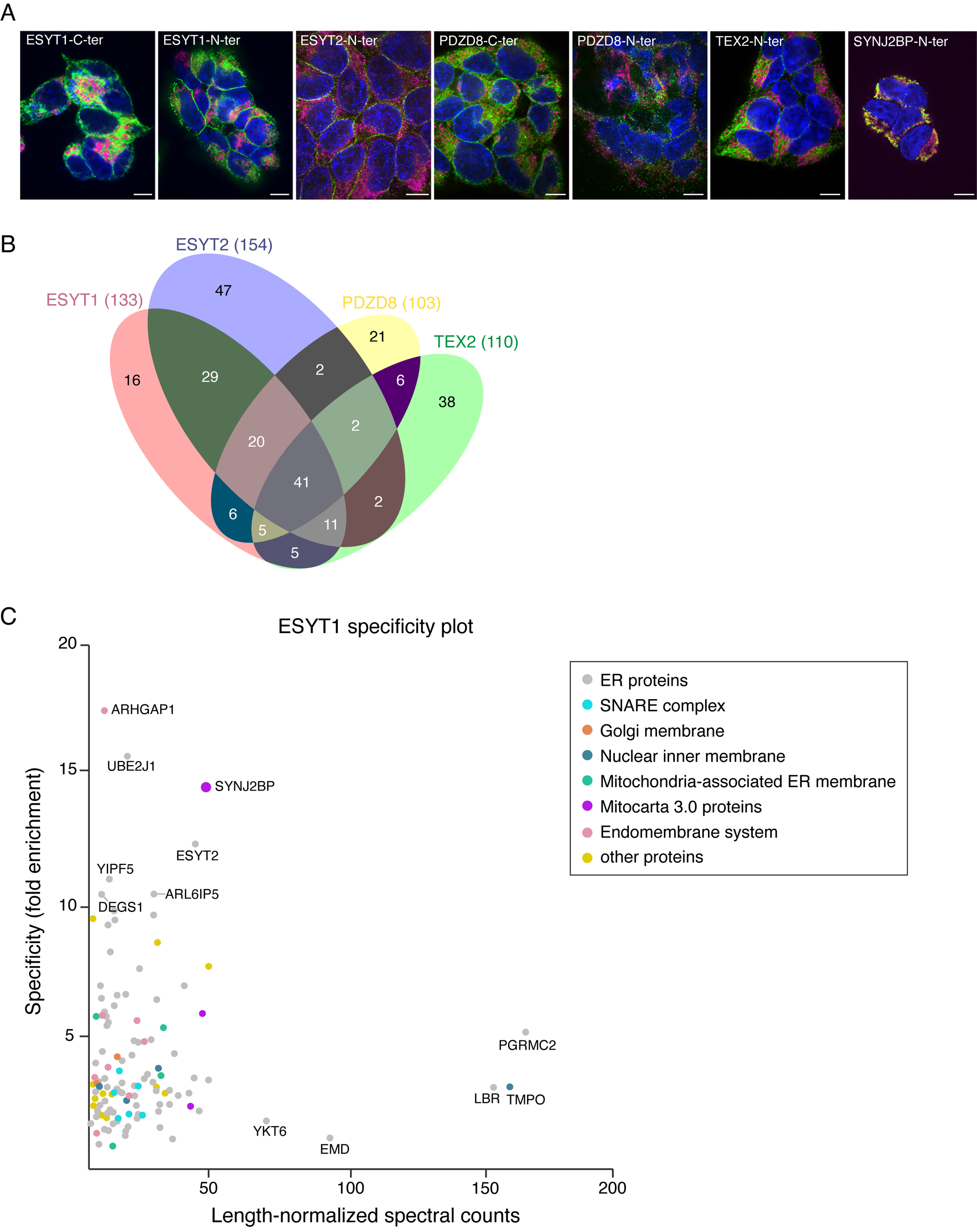

### Figure S2

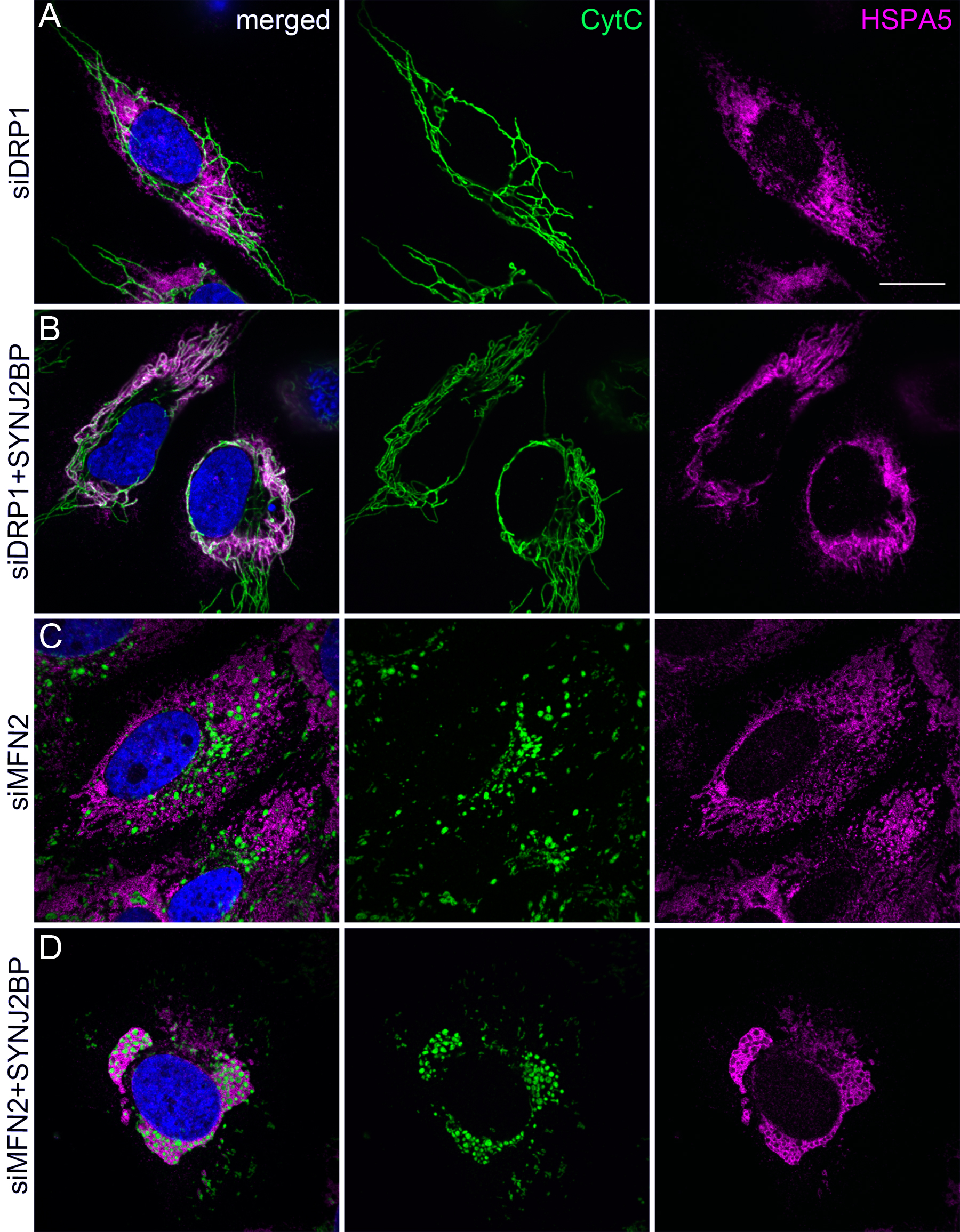

### Figure S3

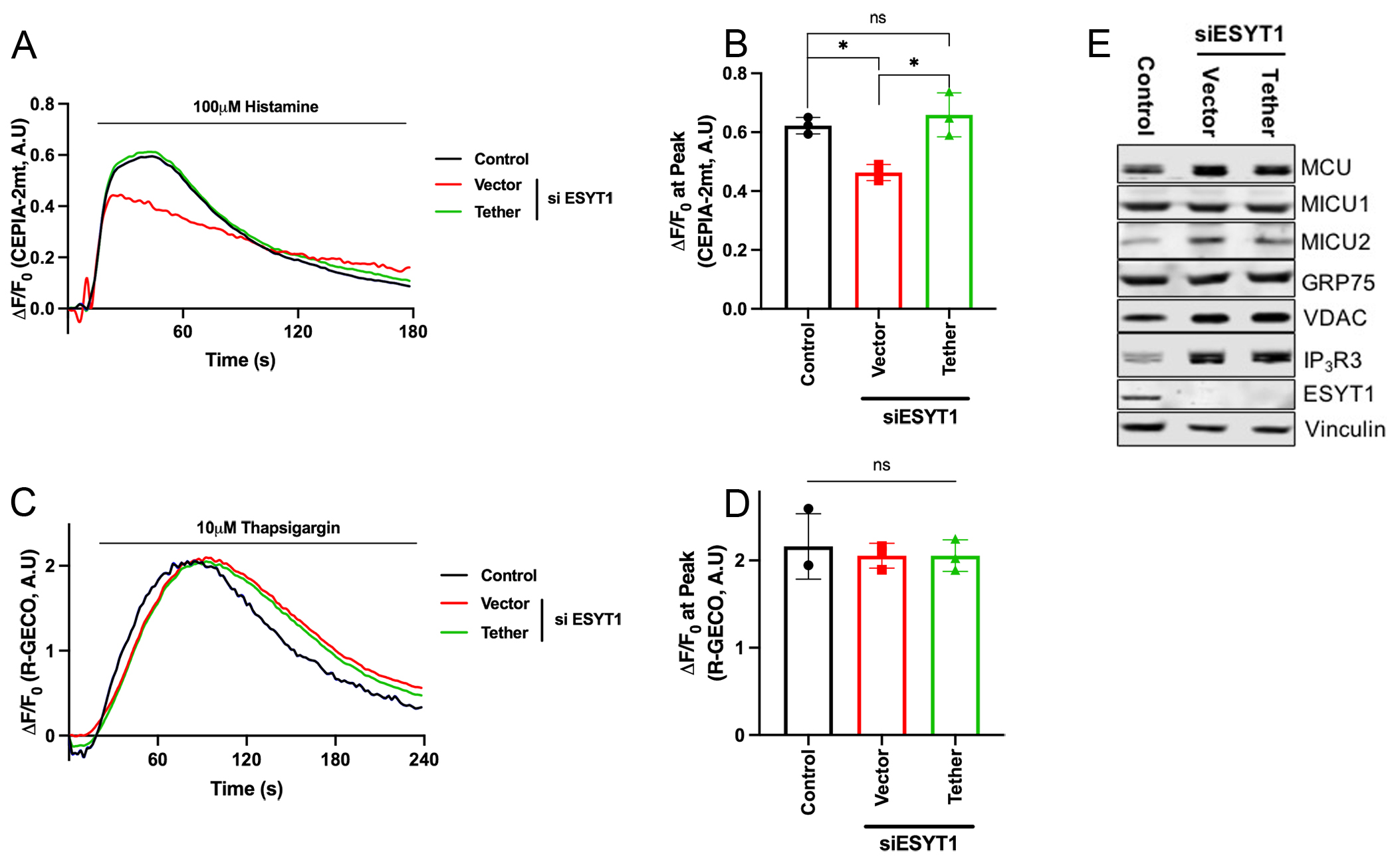

### Figure S4

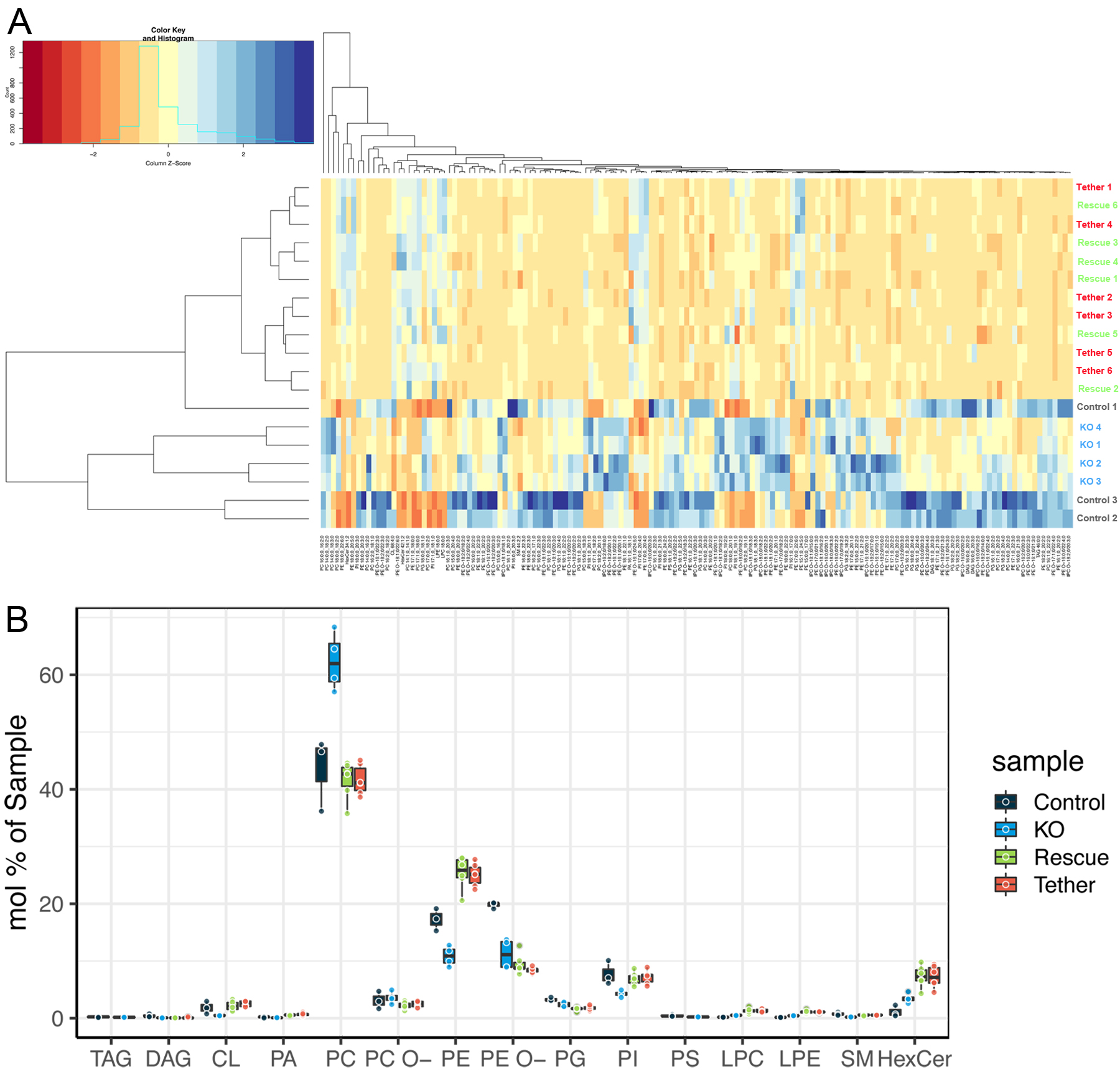

### Figure S5

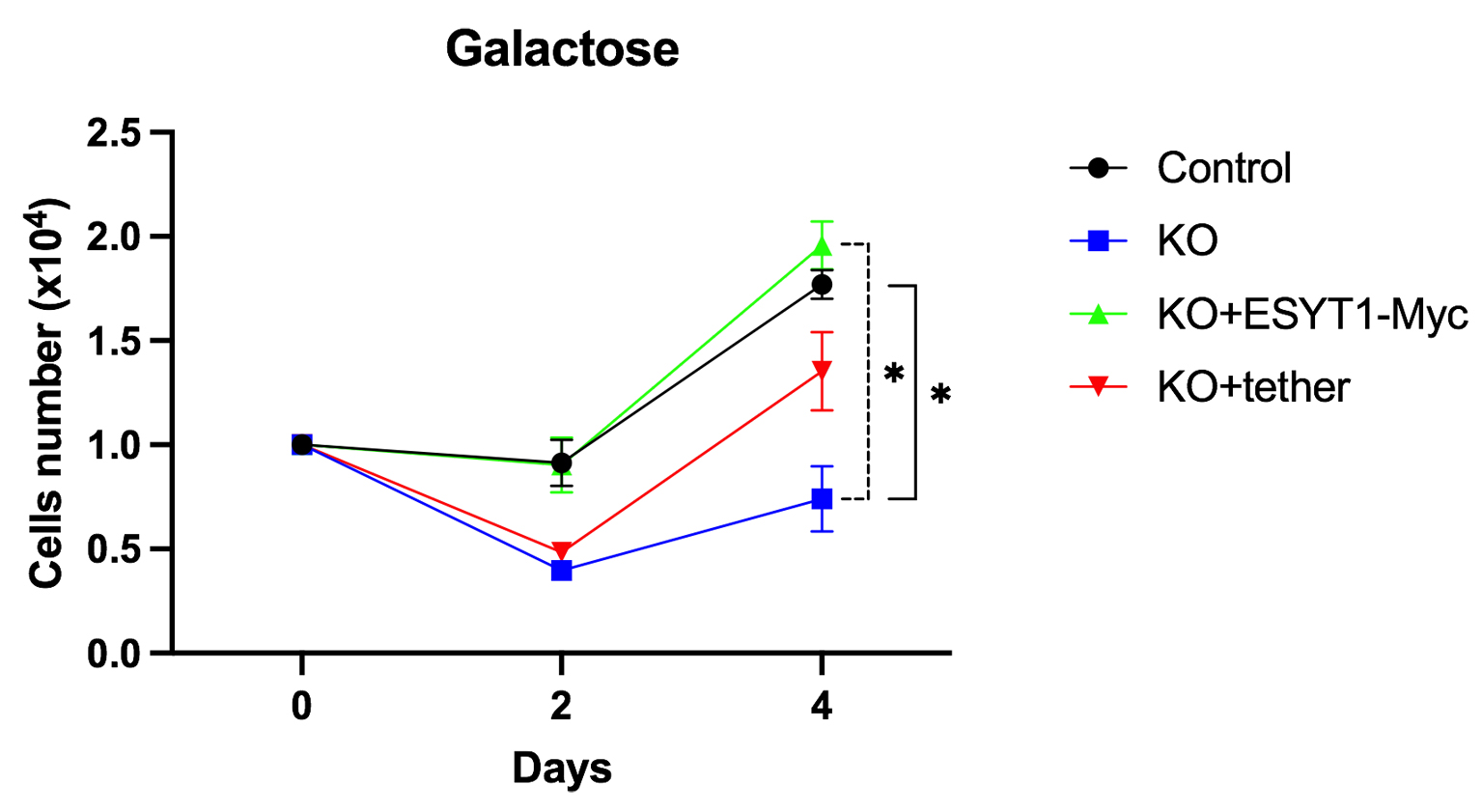
